# Towards autonomous biology: Compiler-Verified Protocols as a Foundation for Real-World AI Execution

**DOI:** 10.64898/2026.05.05.720956

**Authors:** Renjian Song, Yaokai Fu, Ziyan Zhao, Jigang Yu, Qing Yuan, Chang-Ting Chen

**Author notes:** These authors contributed equally to this manuscript (Renjian Song, Yaokai Fu).

## Abstract

Artificial intelligence has advanced from analyzing experimental data to autonomously generating hypotheses, designing experiments, and coordinating closed loop discovery. Yet the translation from computational reasoning to physical execution remains bottlenecked by the experimental protocol, which in biology still relies on ambiguous natural-language descriptions: a medium other engineering disciplines abandoned decades ago in favor of compiler verified specification languages. This deficit fragments reproducibility along three axes: protocol accuracy, pre execution verification, and cross platform portability. Existing formalisms address only subsets of these challenges, trading expressiveness for rigor, portability for standardization, or usability for provenance. Here we introduce the Biology Protocol Language (BPL), a domain specific language with a biology-native type system in which every quantity carries physical units, every reagent declares its physical form, and every container maintains compiler-tracked state, so that implicit assumptions must be stated explicitly and physically impossible operations are rejected at compile time. We further develop BPL-COGEN, a pipeline that couples a fine tuned 30 billion parameter language model with the deterministic compiler in a closed generate validate repair loop, iteratively correcting the translation from natural language SOPs to BPL through compiler diagnostics until all physical, dimensional, and state constraints are satisfied. On a benchmark of 300 published *Nature Protocols* papers, BPL COGEN achieved an overall fidelity score of 95.1 against the source protocols as ground truth. Wet-lab experiment and cross-platform validation in GFP expression library construction and HPLC to UHPLC method translation confirmed that a single BPL source yielded reproducible execution across manual and liquid handler assisted contexts. The results established a novel pipeline that generates compiler-verified protocols, which is an essential prerequisite for physically embodied AI in biology.

## Introduction

Artificial intelligence is rapidly advancing from a tool for analyzing experimental data to an autonomous reasoning engine capable of generating hypotheses, designing experiments, and interpreting outcomes (Boiko et al., 2023). Large language models now propose novel synthetic routes (Bran et al., 2024), multi-agent systems coordinate iterative cycles of prediction and validation (Lu et al., 2024), and self-driving laboratories execute closed-loop optimization across material science and chemistry (Abolhasani & Kumacheva, 2023). Yet a persistent bottleneck separates computational reasoning from physical laboratory reality: the experimental protocol. However sophisticated the upstream AI, its output must ultimately be translated into a precise sequence of physical operations carried out by a human operator, a robotic platform, or both. Today, especially in biology, that translation still relies on natural-language text, a medium that other disciplines abandoned decades ago in favor of formally specified, compiler-verified description languages such as Verilog and VHDL for semiconductor design (Thomas & Moorby, 2002), typed programming languages and deterministic build systems for software engineering (Aho et al., 2006). Biology lacks an equivalent, and this absence is now the rate-limiting step connecting AI-driven experimental design to reproducible physical execution. The deficit extends beyond any single laboratory since science is fundamentally cumulative. Without a shared protocol format, methods cannot be faithfully transferred across sites, platforms, or research groups and thus fragmenting the collaborative infrastructure on which large-scale AI-driven discovery depends.

The consequences of this gap are well documented. A survey of 1,576 researchers found that more than 70% had failed to reproduce another scientist’s experiments and more than half had failed to reproduce their own (Baker, 2016). Standard operating procedures and published methods—the primary vehicles for encoding laboratory knowledge—suffer from ambiguous quantities, inconsistent terminology, implicit assumptions, and unstated execution dependencies (Giraldo, O et al., 2018). These deficiencies compound along three axes. The first is protocol accuracy. A representative instruction from a published transformation protocol—”Resuspend the cell pellet in ice-cold CaCl2 solution, incubate on ice, then plate an appropriate volume onto selective agar”—leaves unresolved the resuspension volume and technique, the CaCl2 concentration, the incubation duration, the meaning of “appropriate volume,” and the identity of the selective agent. Each unstated parameter is a silent branch point; across the dozens of such instructions in a typical protocol, compounding ambiguity makes reproducibility a function of tacit knowledge rather than specification. The second is protocol verification. Natural-language protocols carry no mechanism for checking physical consistency before execution. Protocol inconsistency or fundamental logic errors still rely on manual judgement or even experiment failure to be caught. The third is cross-platform portability. Protocols are written for specific execution contexts, such as equipment. A workflow calibrated for one lab may fail on another due to poor protocol adjustment for equipment availability across labs. A multi-site replication study in synthetic biology found that nominally identical protocols produced two-fold variation in transformation efficiency across four laboratories, driven by unstated differences in execution context rather than biological variability (Beal et al., 2016; Beal et al., 2020).

Over the past two decades, several formalisms have attempted to bridge this gap, yet each addresses only a subset of these challenges. BioCoder encoded protocols as C++ library calls, achieving unambiguous syntax at the cost of expressiveness beyond simple linear workflows (Ananthanarayanan & Thies, 2010). Autoprotocol serialized protocols as parameterised JSON for cloud laboratories but forbids branching logic, precluding conditional workflows (Bates et al., 2017). Antha offered high-level experiment design with automatic reagent computation but imposed proprietary vendor lock-in (Synthace, 2015; Miles & Lee, 2018). LAP coupled standardised scripts to the OpenTrons OT-2, achieving single-platform reproducibility at the expense of portability and compile-time verification (Odriozola et al., 2023). LabOP grounded protocol semantics in RDF/OWL ontologies linked to SBOL, achieving FAIR-compliant provenance at an authoring complexity prohibitive for most bench scientists (Bartley et al., 2023). Oops modelled protocols as probabilistic programs for Bayesian fault diagnosis but sacrificed the granular physical detail needed for direct hardware execution (Shang, M. et al., 2025). No existing solution simultaneously provides compile-time physical verification, hardware-agnostic portability, human–machine interoperability, and a scalable pathway from natural-language SOPs to executable protocols.

To address these gaps, we designed the Biology Protocol Language (BPL) and its generation pipeline Biology Protocol Language Compiler-Guided Generation (BPL-COGEN) (Fig. 1), which together target the three challenges identified above. BPL replaces natural-language ambiguity with formal specification: a biology-native type system in which every quantity carries physical units, every reagent declares its physical form, and every container maintains compiler-tracked states so that what natural language leaves implicit must be stated explicitly, and what is physically impossible is rejected at compile time. BPL-COGEN operationalizes this translation by coupling a fine-tuned 30-billion-parameter language model with a deterministic compiler in a closed generate–validate–repair loop: natural-language SOPs are normalized against a local resource catalogue, converted to BPL, and iteratively corrected against compiler diagnostics until all physical, dimensional, and state constraints are satisfied. On 300 published Nature Protocols papers, BPL-COGEN preserved the scientific content of source *Nature Protocols* articles with 95.1 ± 8.3% overall fidelity while converting heterogeneous inputs into a structurally deterministic, protocol-like representation.

**Fig. 1.**
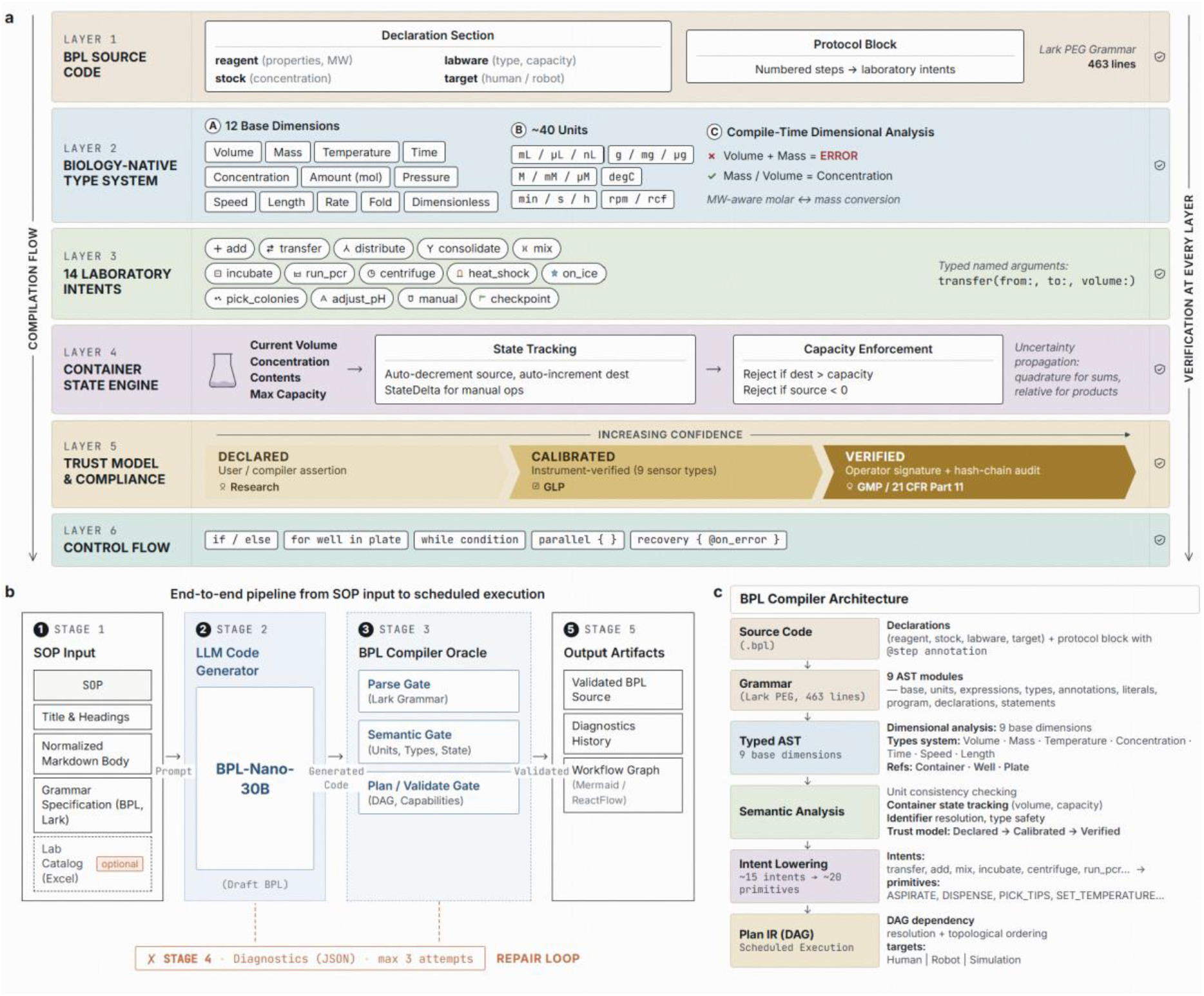
Biology Protocol Language and BPL-COGEN pipeline architecture. **(a)** BPL language architecture. Six-layer structural overview of the Biology Protocol Language. Layer 1: BPL source code comprises a declaration section (reagents, stocks, labware, execution targets) and numbered protocol blocks encoding laboratory intents. The grammar is specified in 463 lines of Lark PEG notation. Layer 2: A biology-native type system enforces compile-time dimensional analysis across 12 base dimensions and approximately 40 laboratory units, rejecting physically impossible operations (e.g., adding a volume to a mass) before execution. Layer 3: Fourteen high-level laboratory intents (transfer, incubate, run_pcr, etc.) accept typed named arguments and abstract over execution details. Layer 4: A container state engine tracks current volume, concentration, contents and maximum capacity for every vessel, automatically updating state on each operation and rejecting programs that would overflow a container or draw from an empty source. Layer 5: A three-tier trust model (Declared, Calibrated, Verified) tracks the provenance and confidence of every state value, supporting Research, GLP and GMP (21 CFR Part 11) compliance contexts. Layer 6: Control flow constructs include conditionals, iteration, parallel execution blocks and structured error recovery handlers. **(b)** Bounded agentic workflow: (1) SOP input with optional lab catalog, (2) BPL-Nano-30B code generation, (3) three-gate compiler validation (parse, semantic, and plan/validate gates), (4) diagnostic-guided repair loop (max 3 attempts), and (5) output artifact generation. **(c)** BPL compiler architecture: six-layer compilation pipeline from source code (declarations and protocol blocks) through grammar parsing (Lark PEG, 463 lines, 9 AST modules), typed AST with dimensional analysis (9 base dimensions), semantic analysis with unit checking and container state tracking (trust model: Declared −> Calibrated −> Verified), intent lowering (~15 intents to ~20 execution primitives), to Plan IR with DAG scheduling for multi-target execution (human, robot, simulation).

To validate actionability and cross-platform portability of protocols generated from BPL-COGEN, we executed compiler-verified protocols in molecular biology (GFP expression library construction) and analytical chemistry (HPLC-to-UHPLC method translation). In the case of GFP library construction, BPL-COGEN compiled two distinct execution contexts—manual and liquid handler assisted—and yielded reproducible outcomes. While for HPLC-to-UHPLC method translation, comparable analytical outcomes of carotenoids characterization were obtained to verify BPL-COGEN’s capability in transitioning protocols based on equipment context.

Overall, the current research establishes a domain-specific language BPL that resolves protocol ambiguity through biology-native types, physical-form constraints, and compiler-tracked container state. Furthermore, the BPL-COGEN pipeline resolves verification deficiency by coupling LLM generation with deterministic compiler validation. BPL-COGEN provides the essential foundation for physically embodied AI in biology by ensuring compiler-verified protocol representations, which is a prerequisite for autonomous laboratory agents.

## Results

### BPL-COGEN provides a standardized protocol generation pipeline

Despite being the most widely used format, Natural-language inherited several problems that prevented it from becoming an accurate format of experiment SOPs. For example, their ambiguity, implicit assumptions, and lack of dimensional typing make them unsafe as direct input to any execution system. To ensure protocol accuracy at the language level, we developed the Biology Protocol Language (BPL) (See Method), a domain-specific language that expresses laboratory protocols as typed, physically constrained programs rather than prose instructions (Fig. 1a).

BPL is organized in six layers, each enforcing a progressively deeper class of constraint (Fig. 1a). The declaration layer defines reagents, concentrations, labware, and execution targets with explicit physical properties. Protocol blocks encode experimental workflows as high-level intents—transfer, mix, incubate, and eleven others—that mirror standard laboratory vocabulary while remaining formally executable. A biology-native type system with compile-time dimensional analysis across nine base physical dimensions and approximately 40 unit annotations ensures that all quantities are unit-consistent and rejects physically invalid operations (e.g., adding a volume of a solid, summing mass and volume) before execution. An intent-lowering layer compiles high-level intents into platform-specific execution primitives, enabling a single program to target human-readable step sheets, robotic liquid handlers, or simulation backends without source modification. A stateful container model tracks material flow automatically—volume, contents, temperature, phase—and enforces constraint checks that prevent infeasible operations such as overfilling or depletion; explicit state-update declarations maintain consistency across non-automated steps. Finally, a compliance layer supports traceability through a three-tier trust model (Declared, Calibrated, Verified) with operator attribution and audit trails for regulated environments.

The language and compiler were iteratively refined through 14 revision cycles, validated against 150 published protocols, and supported by 1,175 test cases spanning dimensional analysis, state tracking, intent lowering, and error recovery, establishing BPL as a robust abstraction for reproducible, cross-platform laboratory execution (See Method).

To make BPL practically useful for scientists without programming backgrounds, a pipeline including BPL compiler oracle and natural language input front end was constructed. The pipeline named Biology Protocol Language Compiler-Guided Generation (BPL-COGEN) converts natural-language SOPs into BPL source code that accurately express protocol intent, using compiler-verified BPL as an intermediate representation. The pipeline operates in five stages: input normalization and resource grounding, grammar-constrained code generation, three-gate compiler validation, diagnostic-guided repair, and deterministic artefact production (Fig. 1b).

Raw SOPs are ingested as Markdown and normalized—canonicalizing whitespace, line endings, and paragraph structure—with title and section metadata extracted to expose the protocol’s logical organization. A laboratory resource catalogue, where available, resolves institution-specific reagent and equipment names against local inventory through bilingual alias matching and relevance ranking, ensuring that generated declarations correspond to physically available materials.

The normalized SOP, complete BPL grammar (463 lines in Lark PEG notation), section structure, and catalogue context form a composite prompt submitted to BPL-Nano-30B (See Method), a fine-tuned 30-billion-parameter language model (see Methods). The grammar includes verbatim; summarizing it substantially reduced first-attempt parse success, as the model required exact production rules to generate syntactically valid code.

Syntactically valid code is not necessarily physically correct—a program may parse yet reference undeclared containers, violate unit consistency, or assume nonexistent hardware capabilities. A deterministic compiler therefore validates each candidate through three sequential gates: the parse gate verifies grammar conformance and produces a typed abstract syntax tree; the semantic gate checks unit consistency, type safety, state coherence, and identifier resolution; and the plan gate performs intent lowering to a directed acyclic graph of execution primitives, dependency-aware scheduling, and backend capability checking. Each gate emits typed diagnostics with error codes, source locations, and targeted repair hints (Supplementary Table 1). When any gate fails, diagnostics are returned to the model as contextualized repair prompts; the corrected program re-enters the compiler, repeating until compilation succeeds or three attempts are exhausted.

Successful compilation emits five deterministic artefacts: (i) the validated BPL source **code** as a version-controllable canonical form; (ii) an execution plan Direct Acyclic Graph (DAG) with resolved volumes, well addresses, and dependency edges; (iii) a diagnostic report with volume-tracking and unit-conversion audit trails; (iv) a workflow graph for visual verification; and (v) a per-step state trace recording container volume, concentration, and trust-level transitions. Together, these artefacts serve as the verified interface between scientific intent and physical execution.

### BPL-COGEN enforces higher protocol accuracy

To quantify the accuracy of BPL-COGEN-generated protocols, we benchmarked the pipeline on 300 published Nature Protocols papers spanning molecular biology, cell culture, biochemistry and analytical chemistry. For each paper, the original natural-language protocol served as the reference description of the experimental procedure. Ten BPL program variants were generated independently per protocol to emulate stylistic variation introduced by different authors. After filtering, this benchmark yielded 2,992 valid processed variants for downstream analysis (see Methods, Benchmark evaluation). Each processed variant was evaluated by a general-purpose LLM along three complementary axes on a 0–100 scale: experiment match, which measures fidelity to the scientific content of the source protocol; protocol validity, which measures whether the output forms a coherent and executable stepwise protocol; and phase completeness, which measures whether all experimentally distinct phases are preserved. The overall fidelity score was defined as the unweighted mean of these three axes.

BPL-COGEN achieved high fidelity across the benchmark, with an overall score of 95.1 ± 8.3 (median 97.0; Fig. 2a). Experiment match, which measures fidelity to the scientific content of the source protocol, remained high at 95.0 ± 9.2. Protocol validity, which measures whether the generated output forms a coherent and executable stepwise protocol, was the strongest axis (98.7 ± 7.6), consistent with the compiler enforcing structurally well-formed outputs. Phase completeness, which measures whether all experimentally distinct stages are preserved, was slightly lower at 91.0 ± 10.4, mainly reflecting cases in which ambiguously bounded phases in the source text were merged during normalization rather than omitted altogether.

**Fig. 2.**
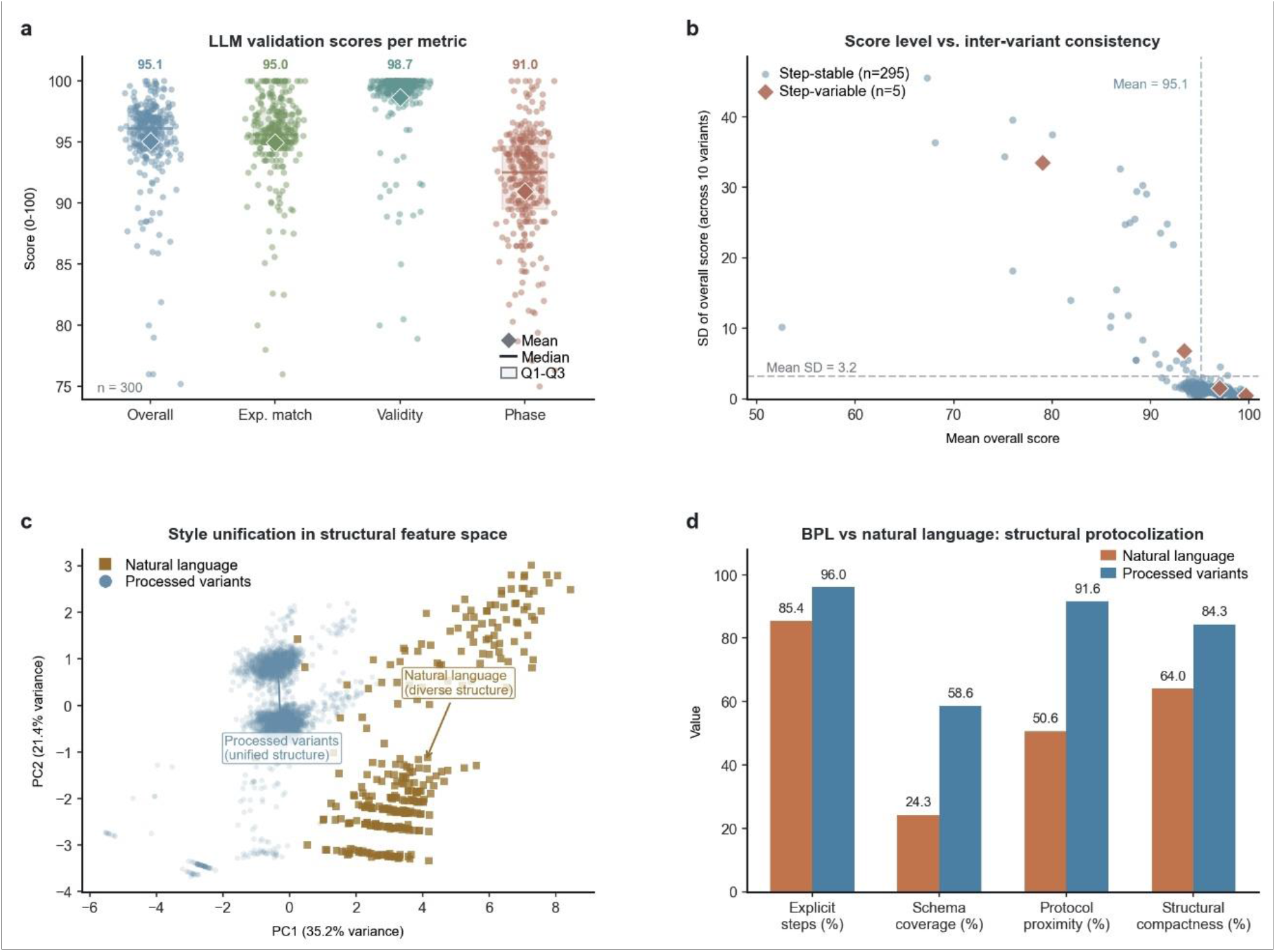
LLM-based validation of BPL protocol generation across Nature Protocols benchmarks. **(a)** LLM validation scores across processed variants. Overall score, experiment match, protocol validity and phase completeness all remained high (means: 95.1, 95.0, 98.7 and 91.0, respectively; n = 300 papers, 2,992 variants). This panel establishes that BPL outputs are not only structurally regularized, but also remain faithful to the underlying experiment. **(b)** Inter-variant consistency at the paper level. Each point represents one source paper, positioned by its mean overall score across 10 variants and the standard deviation of that score across variants. Blue points denote step-stable papers and red diamonds denote step-variable papers. Most papers cluster in the high-score, low-variance regime; 295 of 300 papers are step-stable whereas only 5 are step-variable. **(c)** Principal component analysis of structural document features for 300 raw natural-language source papers and 2992 processed variants. PCA was performed on six document-level structural features: word count, line count, step count, bold-format count, CRITICAL-token count and dash-bullet count. Raw source papers occupy a diffuse region of feature space, whereas processed variants collapse into a much tighter protocol-like manifold. **(d)** Quantitative summary of structural protocolization before and after BPL. Explicit step structure denotes the proportion of documents containing at least one detected procedural step; schema coverage denotes the proportion containing at least three canonical protocol markers; protocol proximity quantifies how close documents lie to the centroid of processed-protocol space in PCA coordinates; and structural compactness quantifies how tightly documents cluster around their own class centroid in the same space. Relative to raw source papers, processed variants show higher rates of explicit step structure (85.4% −> 96.0%), broader canonical protocol schema coverage (24.3% −> 58.6%), greater proximity to the processed-protocol region in PCA space (50.6% −> 91.6%) and higher structural compactness (64.0% −> 84.3%)

We next asked whether independently generated variants of the same source paper converged after BPL processing. Cross-variant consistency was high: 295 of 300 papers (98.3%) were step-stable across all 10 independent runs, clustering in a high-score, low-variance regime (mean s.d. = 3.2; Fig. 2b). The five step-variable cases involved source protocols with ambiguous procedural structure, although all retained overall scores above 80. These results indicate that BPL not only preserves protocol fidelity, but also drives multiple rewrites of the same experiment toward a common processed representation.

To test whether this convergence reflected a broader structural transformation, we projected documents into a shared structural feature space using PCA of six document-level features: word count, line count, step count, bold-format count, CRITICAL-token count and dash-bullet count. These features capture document length, explicit procedural segmentation, emphasis markers and list-like formatting, and therefore summarize how strongly a document resembles an explicit protocol rather than a free-form narrative description. In this space, raw source papers were broadly dispersed, whereas processed variants collapsed into a much tighter protocol-like manifold (Fig. 2c), indicating corpus-level structural unification rather than merely paper-specific convergence.

This transformation was also evident in direct structural metrics. In Fig. 2d, explicit steps refers to the fraction of documents containing at least one clearly detected procedural step, and therefore captures whether actions are written in an explicit stepwise form. Schema coverage refers to the fraction of documents containing at least three canonical protocol markers, such as objective, materials, expected results, troubleshooting, critical-step language or explicit step enumeration, and therefore measures how completely standard protocol sections are surfaced. Protocol proximity measures how close documents lie to the centroid of processed-protocol space in PCA coordinates, and therefore reflects how strongly a document resembles the overall structural profile of BPL-standardized protocols. Structural compactness measures how tightly documents cluster around their own class centroid in the same space, and therefore reflects within-class structural uniformity. Relative to raw source papers, processed variants showed higher rates of explicit step structure (85.4% −> 96.0%), broader canonical protocol schema coverage (24.3% −> 58.6%), greater proximity to the processed-protocol region in PCA space (50.6% −> 91.6%) and higher structural compactness (64.0% −> 84.3%; Fig. 2d). Together, these results define a clear progression: BPL outputs retain the scientific content of the original protocol, independently generated variants converge toward the same processed form, and the resulting documents adopt a more uniform protocol-like structure. This combination of semantic fidelity and structural reproducibility is essential for robotic execution, protocol comparison and regulatory archiving.

### Pre-Execution Protocol Verified Through Compiler-Driven Diagnostics

Natural-language protocols carry no intrinsic mechanism for checking physical consistency before execution. Dimension mismatches, capacity overflows, and undeclared reagents are typically caught only through manual review or, worse, through failed experiments at the bench — consuming reagents, instrument time, and operator effort before the error is ever surfaced. To address this limitation, BPL-COGEN enforces protocol correctness via compile diagnostic: candidate programs must pass three sequential validation gates (parse, semantic, plan) before any physical resources are committed (Fig. 1b, stage 4). At each gate, the compiler emits structured diagnostics containing an error code, severity, source location, and a targeted repair hint, transforming protocol review from post-hoc troubleshooting into pre-execution verification. We applied this compiler diagnostics through a benchmark corpus to evaluate the efficiency of the self-correction.

In this evaluation, the compiler issued 343 diagnostics, distributed across five categories (Fig. 3a, Supplementary Table 1). Dimension mismatches — a value of one physical quantity assigned to a parameter expecting another (for example, a mass value given where a volume is required) — were the most frequent class (142 occurrences, 41.4%). Capacity violations, in which a transfer would exceed a container’s declared volume, accounted for 87 diagnostics (25.4%). Undeclared identifiers, where reagents or labware referenced in the procedure were absent from declarations, contributed 64 (18.7%). State conflicts — operations that produce physically impossible conditions such as negative volumes — accounted for 38 (11.1%). Trust violations, arising when a GLP or GMP protocol references values without the required calibration provenance, were least frequent (12, 3.5%). Fig. 3c illustrates a representative dimension-mismatch case: a transfer operation specifying volume: 50 mg — a mass value assigned to a volume parameter — is rejected by the compiler, which emits a JSON diagnostic pinpointing the source location (line 11, column 13) and suggesting the corrective unit class (“Use mL, uL, or nL”). Upon receiving this diagnostic, the language model replaces the erroneous argument with volume: 50 uL, and the corrected program passes all three validation gates.

**Fig. 3.**
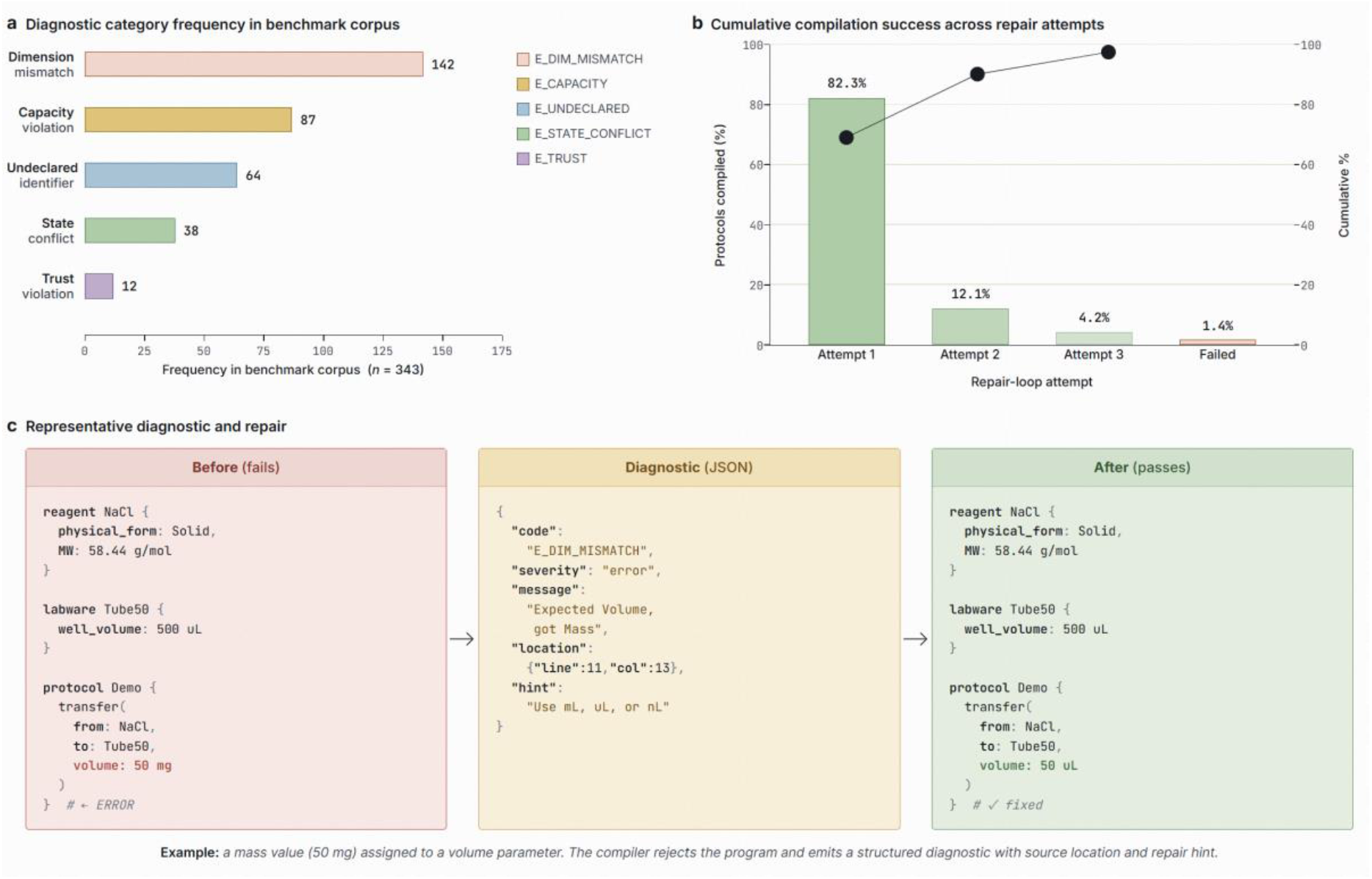
Compiler-driven diagnostics and repair loop performance. **(a)** Frequency of diagnostic categories across the benchmark corpus. Dimension mismatches are the most common failure mode (142 occurrences), followed by capacity violations (87), undeclared identifiers (64), state conflicts (38), and trust violations (12). **(b)** Compilation success rate by attempt number: 82.3% of protocols compile on the first attempt; the repair loop resolves an additional 16.3%, leaving only 1.4% unresolved after three attempts. **(c)** Representative before-and-after example: a mass value (50 mg) assigned to a volume parameter is caught by the compiler with a structured diagnostic (JSON) containing the error code, source location, and targeted repair hint.

Critically, the compiler diagnostics also enabled automated self-correction by the language model. BPL-COGEN achieved 82.3% first-attempt compilation success on the benchmark corpus after fine-tuning (Fig. 3b). Of the variants that required repair, 12.1% compiled after one diagnostic-guided iteration and 4.2% after two, yielding a cumulative compilation rate of 98.6% within two iterative loops. Only 1.4% of variants remained unresolved, typically involving source SOPs with deeply ambiguous or contradictory specifications that could not be resolved without human clarification. The high repair rate confirms that source-localized repair hints provide sufficient signal for the language model to correct its own errors without human intervention in the vast majority of cases — a closed-loop verification capability that has no analogue in conventional natural-language protocol workflows.

### Cross-Platform Protocol Portability: From Molecular Biology to Analytical Chemistry

One key obstacle BPL-COGEN aimed to address was cross-platform portability. Ideally, a single source program can compile to respective execution protocols based on lab specific context such as equipment availability or execution preferences. To validate this feature, we executed compiler-verified protocols in two independent domains: molecular biology (GFP expression library construction) and analytical chemistry (HPLC-to-UHPLC method translation).

### Validation of manual and liquid handler assisted protocols generated by BPL-COGEN

To validate the actionability of BPL-COGEN-generated protocols, a case study was conducted focusing on the construction of a GFP plasmid library in *E*.*coli*. BPL-COGEN was prompted to design 11 distinct expression plasmids (pBEE_TrcT7_DasherGFP, pBEE_Trcp1_DasherGFP through pBEE_Trcp10_DasherGFP) utilizing a shared vector backbone and a Dasher GFP insert (Duran et al., 2013), driven by 11 unique promoters (TrcT7, Trcp1 through Trcp10, Supplementary Table 2). To test BPL-COGEN’s ability to generate both human-readable instructions for manual operation and machine-readable worklists for liquid handler assisted execution on liquid handlers, two corresponding protocols were generated for polymerase chain reaction (PCR), DNA assembly, and transformation (Fig. 4a). Both the manual protocol (supplementary file 1) and the Biomek protocol and worklists (supplementary file 2 - 8) were generated from a single BPL source program without any source modification, demonstrating the language’s flexibility between execution modes.

**Fig. 4.**
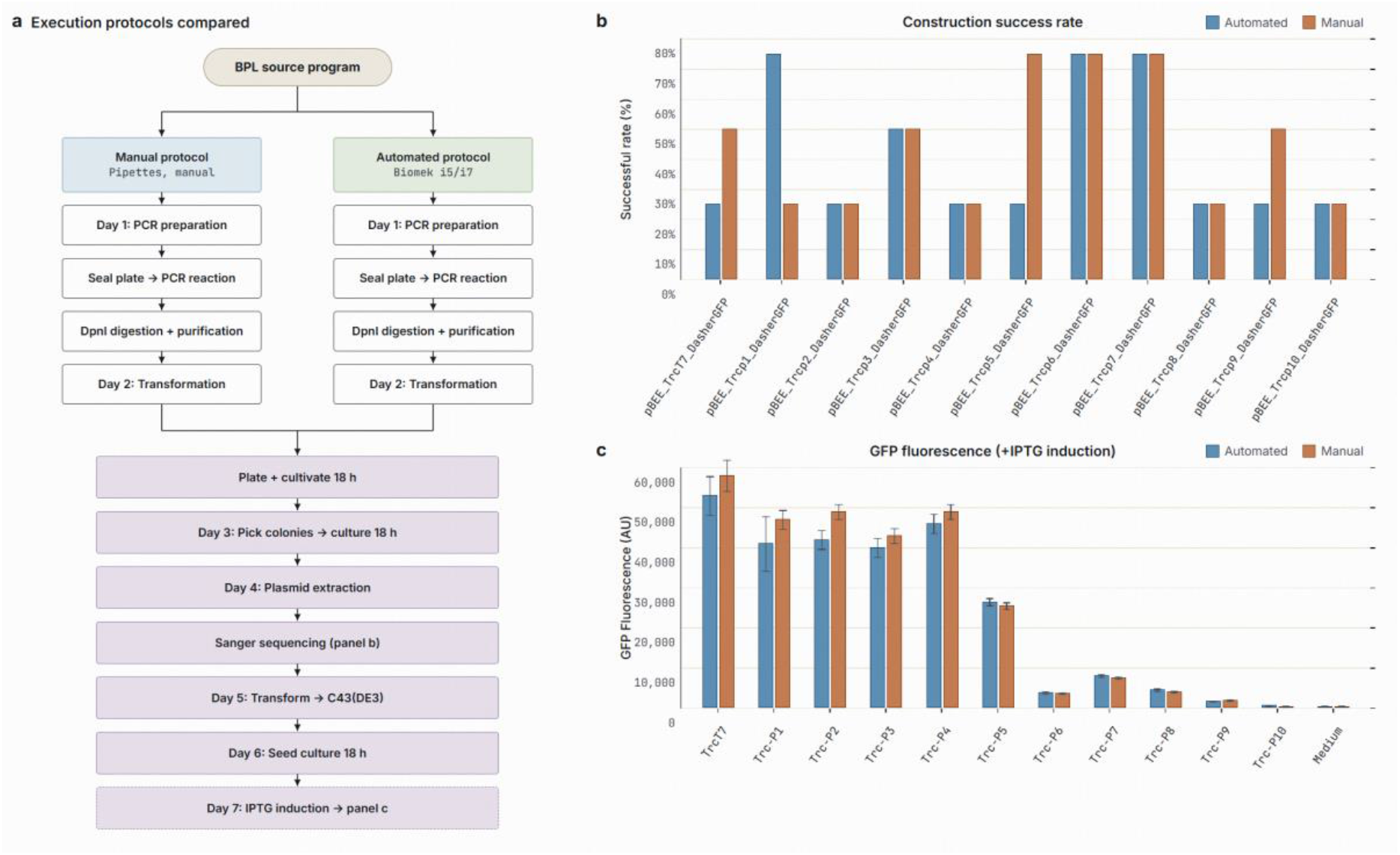
Wet-lab validation of BPL-COGEN protocols. **(a)** Overview of the experiment workflow. To validate both manually operated and liquid handler–based protocols, BPL-COGEN generated corresponding instructions for PCR, DNA fragment assembly, transformation, plating, and colony picking/screening procedures. **(b)** Construction success rate of GFP-expressing plasmids using manual or liquid handler assisted (automated) protocols. For each plasmid, 4 colonies were picked for Sanger sequencing. **(c)** GFP fluorescence measurement of cell cultures with IPTG induction. Medium: negative control without cells.

Gel electrophoresis of the PCR products demonstrated comparable results across both the manual and liquid handler assisted, with all target DNA fragments successfully amplified (data not shown). Following the Gibson assembly (Gibson et al., 2009) of the amplified DNA fragments, the respective plasmids were transformed into DH5a competent cells. Subsequent sequencing results confirmed a similar construction success rate using the liquid handler assisted and manual workflows (Fig. 4b), indicating that BPL-COGEN generates reliable protocols for both manual and machine-executable workflows.

To further evaluate the phenotypic variance across the promoter library, the verified plasmids were then transformed into *E. coli* C43(DE3) and cultivated for fluorescence measurement. As demonstrated, the 11 corresponding strains displayed a clear gradient of fluorescence intensity upon isopropyl beta-D-1-thiogalactopyranoside (IPTG) induction (Fig. 4c). Crucially, this distinct expression gradient was consistently observed across both the manually handled and liquid handler assisted sample sets. Altogether, these results demonstrate that BPL-COGEN-generated experimental protocols yield highly comparable and reproducible results across both human-executed and robot-executed biological pipelines, and that the protocols generated lead to successful and meaningful molecular biology outcomes.

### Protocol Translation Across Different Analytical Instruments Facilitated by BPL-COGEN

To evaluate BPL-COGEN beyond molecular biology, we tested the pipeline on translating a validated HPLC method for five fat-soluble compounds (retinol, retinal, retinyl acetate, lycopene, and β-carotene) to a UHPLC platform with a different column geometry (see Methods). The reference method required 32 min per run on a conventional C18 column (4.6 × 250 mm, 5 μm); the target was a sub-2 μm core-shell column (2.1 × 50 mm, 1.7 μm).

Given the two column specifications and the original gradient program as input, BPL-COGEN applied the scaling formulae of the 2025 Chinese Pharmacopoeia General Chapter 0512 (Chinese Pharmacopoeia Commission, 2025) attached as prompts and generated a complete translated method (Fig. 5a). The compiler-calculated parameters (flow rate 0.613 mL/min, injection volume 0.42 μL, gradient duration 2.18 min) closely matched the values adopted after minor practical rounding (0.6 mL/min, 1 μL, 2.1 min), achieving a 14.7-fold reduction in run time and 95.8% reduction in solvent consumption.

**Fig. 5.**
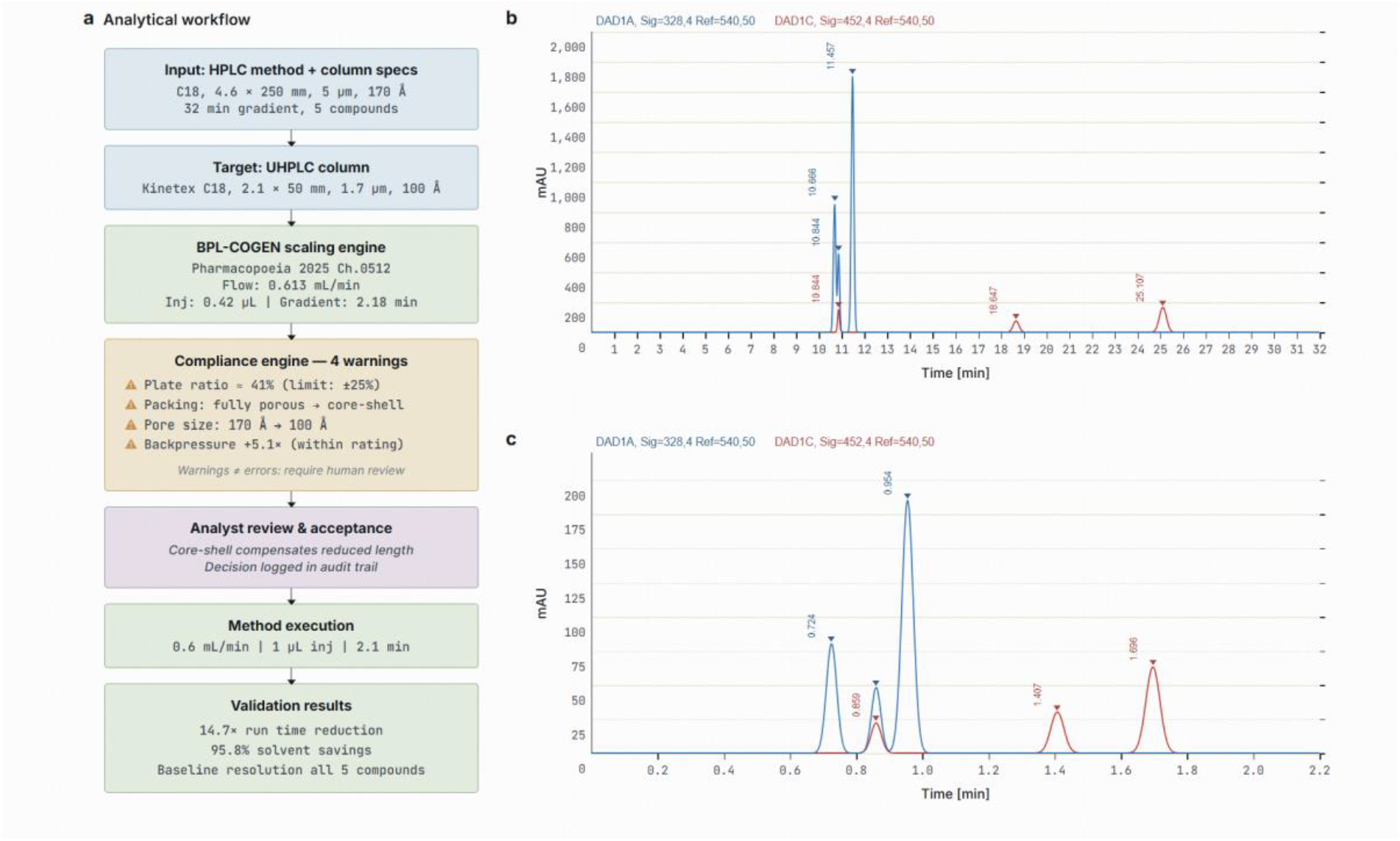
Chromatographic validation of BPL-COGEN-generated HPLC-to-UHPLC method translation. **(a)** Translation workflow. BPL-COGEN scaled the original method using 2025 Chinese Pharmacopoeia formulae (Ch. 0512). The compliance engine flagged four warnings (plate ratio, packing type, pore size, backpressure); the analyst reviewed and accepted all deviations with decisions logged in the audit trail. **(b)** HPLC chromatogram (C18, 4.6 × 250 mm, 5 μm; 32 min). (c) UHPLC chromatogram (Kinetex C18, 2.1 × 50 mm, 1.7 μm; 2.1 min). All five compounds (retinol, retinal, retinyl acetate, lycopene, β-carotene) achieved baseline resolution with preserved elution order.

Before execution, the BPL compliance engine issued four warnings — flagged deviations from pharmacopoeial equivalence thresholds, distinct from compilation errors that halt the pipeline. The warnings identified a 41% reduction in theoretical plate ratio exceeding the 25% equivalence limit, a packing-type change from fully porous to core-shell, a pore-size category change (170 Å to 100 Å), and a 5.1-fold backpressure increase within the column’s rated limit. This distinction between errors and warnings is by design: errors indicate physically invalid operations; warnings flag methodological changes that alter the analytical basis and require informed human judgement. The analyst reviewed all four warnings, accepted the deviations on the basis that core-shell packing compensates for reduced column length through higher per-unit-length efficiency, and the compliance system documented this decision in the audit trail.

The translated method preserved elution order and achieved baseline resolution for all five compounds (Fig. 5b and c), including the critical pair retinol and retinal (Resolution = 1.2), which were separated by only 0.17 min on the original method. Retention times compressed proportionally from the 10–25 min range to 0.72–1.69 min, consistent with the predicted scaling factor. This result demonstrates two capabilities that manual method translation lacks: automated dimensional scaling with full audit trail, eliminating the recalculation errors that are the most common source of method-transfer failures; and compliance-aware verification that ensures deviations from pharmacopoeial standards are explicitly flagged, reviewed, and documented before execution — a GMP requirement that is difficult to enforce with natural-language SOPs.

## Discussion

In this study, we developed BPL-COGEN, a compiler-guided generation pipeline that translates natural-language laboratory protocols into verified, executable programs, and demonstrated 95.1% semantic fidelity across 300 published *Nature Protocols* articles. Underpinning this pipeline is BPL itself — a domain-specific language with compile-time verification of physical and logical consistency, drawing on the formal-verification tradition that has long underwritten reliability in compiler and systems engineering (Leroy, 2009). Together, they bridge the gap between biology’s accumulated natural-language knowledge and the structural rigor that reproducible, automated science demands.

In bridging the persistent gap between computational reasoning and physical execution, BPL-COGEN systematically addressed the three fundamental challenges of natural-language standard operating procedures: protocol accuracy, pre-execution verification, and cross-platform portability. First, BPL established protocol accuracy by replacing ambiguous prose with a biology-native type system in which every quantity is bound to physical units and every container state is tracked, eliminating the implicit assumptions that plague manual protocol replication (Giraldo, O et al., 2018). Second, the pipeline introduced a paradigm shift in protocol verification through its closed-loop, compiler-guided architecture. By routing language-model proposals through deterministic semantic and physical validation gates, BPL-COGEN detected and repaired dimension mismatches, capacity overflows, and logic conflicts prior to any physical execution, achieved a 98.6% compilation success rate within two iterative loops. This pattern — coupling a generative model to a deterministic verifier — extended a broader trend of grounding large language models in external tools and formal oracles (Schick et al., 2023; Gao et al., 2023). Third, BPL-COGEN delivered cross-platform portability by separating high-level experimental intent from hardware-specific execution primitives. As shown by the translation of a single protocol source into both manual bench instructions and liquid-handler worklists for a GFP plasmid library, and by the seamless scaling of an HPLC method to UHPLC, the framework supported reliable execution across variable laboratory environments.

Two independent wet-lab case studies validated BPL-COGEN’s capacity to generate actionable protocols with meaningful scientific outcomes. In the GFP expression library construction, a single BPL source program for the experiment was compiled to a human-readable manual instructions and a machine-executable Biomek i7 worklist. The experiment results demonstrated successful assembly of all designed plasmids via Gibson assembly (Gibson et al., 2009) for both protocols. Verification by sequencing and fluorescence measurement indicated similar outcomes. In the other case study, the BPL-COGEN converted a 32-minute conventional HPLC method to a 2.1-minute UHPLC method with comparable results, illustrating the pipeline’s utility for analytical method adaptation across instrument configurations. Notably, the conversion engine applied pharmacopoeia-compliant scaling formulae (Chinese Pharmacopoeia General Chapter 0512) and generated actionable compliance warnings — flagging a theoretical plate ratio reduction, packing-type change, and backpressure increase — that guided the analyst’s experimental verification before adopting the converted method. The cross-domain validation — spanning constructive molecular biology and analytical measurement — demonstrated BPL-COGEN’s compiler-guided architecture generalizes beyond any single protocol family, provided the domain’s operations can be encoded within BPL’s intent vocabulary and type system. The results not only validated the actionability of the BPL generated protocols, but also demonstrated the key capability of BPL-COGEN on protocol portability, accuracy, and verification.

The structural-determinism finding has implications that extend beyond protocol execution. Version control, automated comparison, and regulatory audit of laboratory procedures all benefit from canonical representations that eliminate the structural noise inherent in natural language. When individual researchers independently describe the same procedure, natural-language descriptions diverge in step count, operation granularity, and implicit assumptions a divergence that underlies the cross-laboratory variability repeatedly documented in synthetic biology (Beal et al., 2016; Beal et al., 2020). BPL enforces a different regime: 98.3% of protocols in our benchmark remained step-stable across ten independent generations, and document-level feature analysis showed that processed variants collapse from the diffuse feature space of raw prose into a tight protocol-like manifold. Combined with the three-tier trust model (Declared → Calibrated → Verified) and hash-chained audit trail, this canonicalization provides a foundation for machine-verifiable GxP compliance and for programmatic operations on the scientific record — protocol diffing, meta-analysis of methods at corpus scale, and machine-readable regulatory dossiers — none of which are feasible over free-form prose.

The compiler-driven diagnostics represent a qualitative shift in protocol validation. Traditional protocol review relies on expert manual inspection and is often reactive, surfacing errors only after failed experiments have consumed reagents, instrument time, and operator effort — a pattern that contributes to the reproducibility crisis in which more than 70% of researchers report failing to reproduce another scientist’s experiments (Baker, 2016). The deterministic compiler instead captures errors at compile time, emitting systematic, structured diagnostics with source locations and targeted repair hints, and — crucially — the same diagnostics close the loop for the language model itself, enabling 82.3% first-attempt success and 98.6% resolution within two iterations without human intervention. This pre-execution verification transforms protocol quality assurance from reactive troubleshooting into proactive error prevention, a closed-loop capability that has no analogue in conventional natural-language protocol workflows.

Several limitations should be noted. Phase completeness (91.0%) was the weakest evaluation axis, reflecting an intent vocabulary optimized for wet-lab operations and less expressive for computational or imaging-heavy protocols; only 2 of 300 protocols (whole-genome sequencing analysis and bacteriophage engineering) averaged below 70%, both involving predominantly computational workflows. Our corpus-level evaluation relied on LLM-as-judge scoring for experiment match and phase completeness; while calibrated against deterministic compiler signals for protocol validity, this approach inherits the known biases of model-based evaluation including position, verbosity, and self-preference biases (Zheng et al., 2023) — and independent human re-scoring of a stratified subsample would further strengthen the claim. Wet-lab validation covered two domains at a single organization, and multi-site replication — the pattern that originally surfaced cross-laboratory variability in synthetic biology (Beal et al., 2016) — remains the decisive test of portability. Finally, the compiler-guided approach presupposes a formal grammar and a constraint oracle; for domains lacking such infrastructure, the agentic repair loop cannot be directly applied, and our own experience — fourteen grammar revisions and 1,175 test cases — indicates that bootstrapping this substrate in a new domain is a non-trivial undertaking.

Ultimately, BPL-COGEN establishes a critical infrastructural layer for the future of reproducible and automated science. By translating heterogeneous natural-language texts into structurally deterministic and semantically faithful executable programs, this work demonstrates that the historical ambiguities of legacy protocols can be computationally resolved without requiring bench scientists to learn a new coding language. Furthermore, the generalizable nature of the compiler-oracle pattern suggests that this approach can extend beyond biology into chemistry, materials science, and clinical protocol formalization, paving the way for a universally interoperable and machine-verifiable scientific ecosystem. The BPL framework that operates as an intent-to-execution translation engine can be integrated with upstream AI-driven hypothesis generators (Boiko et al., 2023; Bran et al., 2024) and downstream robotic execution platforms (Abolhasani & Kumacheva, 2023) to enable fully autonomous, closed-loop discovery systems. Taken together, the reported BPL-COGEN not only mitigated the substantial economic and scientific costs of experimental irreproducibility (Baker, 2016), but also laid the essential groundwork for physically embodied AI in biology by providing a compiler-guided generation paradigm.

## Materials and Methods

### BPL language design

The Biology Protocol Language was designed through an iterative, empirically grounded process that prioritized laboratory domain fidelity over general-purpose expressiveness. The language design proceeded in three phases (Supplementary Fig. S1).

In the first phase (domain analysis), we systematically analyzed 150 published protocols from *Nature Protocols*, JOVE, and institutional SOP repositories, cataloguing the atomic operations, physical quantities, container types, control flow patterns, and compliance requirements that recur across molecular biology, biochemistry, and analytical chemistry workflows. This analysis identified 14 distinct laboratory intents that collectively cover over 95% of operations encountered in the corpus (transfer, add, mix, incubate, centrifuge, run\_pcr, pick\_colonies, adjust\_ph, heat\_shock, on\_ice, distribute, consolidate, manual, and checkpoint), as well as the dimensional categories and unit systems required to express their parameters without ambiguity.

In the second phase (grammar specification), we encoded these domain constraints as a formal grammar using the Lark parsing framework’s PEG notation (Shinan, 2017). Lark was selected for its native support for ambiguity-free parsing, priority-based rule ordering (critical for distinguishing compound units such as mg/mL from division expressions), and direct generation of typed parse trees without a separate lexer stage. The grammar (463 lines, version 2.4) defines a two-section program structure — declarations followed by protocol blocks — that mirrors the natural organization of laboratory protocols: a materials-and-equipment preamble followed by a sequential procedure. Each declaration type (reagent, stock, labware, target) carries mandatory typed attributes that the compiler enforces at parse time, eliminating an entire class of downstream errors. The grammar underwent 14 major revisions driven by conformance testing against the protocol corpus, with each revision expanding coverage (adding parallel execution blocks, recovery handlers, worklist intents for high-throughput operations) while maintaining backward compatibility with previously validated programs.

In the third phase (type system and compiler construction), we implemented the biology-native type system with compile-time dimensional analysis across 12 base dimensions (volume, mass, temperature, time, concentration, amount, pressure, speed, length, rate, fold, and dimensionless), a unit registry of approximately 40 laboratory units with automatic compound dimension derivation, and molecular-weight-aware concentration conversion between mass and molar representations. The compiler was structured as a six-gate pipeline: parse, semantic analysis, plan/DAG construction, experiment-parameter validation, capability checking, and backend preflight. This layered architecture ensures that each gate can be independently tested and extended — the test suite comprises 1,175 tests — and that diagnostic messages precisely identify the gate, source location, and suggested repair for any rejected program.

### Model fine-tuning

#### Training dataset construction

A four-layer synthetic data pipeline (Supplementary Fig. S2) was developed to convert 47 standard operating procedures (SOPs) into paired natural-language-to-BPL training examples.

Layer 1 — Seed expansion. Each SOP was expanded into 5–10 instruction scenarios drawn from a 10 × 4 matrix of ten archetypes and four complexity tiers. The archetypes span standard execution, scale variation, automation adaptation, troubleshooting, teaching, time-constrained, resource-constrained, quality-controlled, GMP-documented, and optimized protocols; the sampling distribution across archetypes was weighted by each SOP’s complexity tier. This step produced ~300–400 expanded scenarios. Layer 2 — BPL generation. Candidate programs were generated for each scenario with Qwen-Plus via the DashScope API and validated by a six-point compiler check: syntactic correctness, unit validity, volume conservation, capacity bounds, complexity compliance, and GMP compliance. Candidates failing any check were regenerated, with up to eight retries per scenario. Layer 3 Quality critique. Surviving candidates were scored by an LLM critic on four axes weighted equally at 25 points each — scientific accuracy, BPL syntax, protocol completeness, and instruction alignment — against a passing threshold of 80/100. Records scoring below threshold entered a self-refinement loop for up to four iterations. Layer 4 — Evolutionary diversification. A random 30% of approved records were mutated along one of five dimensions (scale, error handling, optimization, automation, or quality control) and re-validated through Layers 2–3. Compound filtering across the four layers yielded 2,714 validated instruction– protocol pairs, exported as bpl_train_alpaca.jsonl and bpl_train_full.jsonl.

#### Fine-tuning configuration

The corpus was formatted with the Nemotron chat template (system/user/assistant turns) using response-only loss masking (label = −100 on all non-response tokens). Low-Rank Adaptation (LoRA; Hu et al., 2022) was applied to every linear projection of Nemotron-3-Nano-30B-A3B (NVIDIA, 2025) with rank *r* = 8, α = 16, and dropout = 0.0, yielding 221 M trainable parameters — 0.69 % of the 31.8 B base model.Training ran with SFTTrainer at an effective batch size of 8, peak learning rate 2 × 10^−4^ under linear decay, 5 warmup steps, and 1 epoch (340 optimizer steps). The run completed in 72 min on a single NVIDIA RTX PRO 6000 (94.97 GB VRAM) and reduced training loss from 1.40 to 0.37 (73.4 % reduction).

#### Deployment

The fine-tuned model generates syntactically valid BPL from natural-language instructions with explicit chain-of-thought reasoning. Checkpoints were released in three formats: raw LoRA adapters, fp16 merged weights, and GGUF quantizations (Q4_K_M and Q5_K_M) for inference via llama.cpp or Ollama.

### Benchmark Design

#### Benchmark design

For each source paper in *Nature Protocols*, 10 BPL program variants were generated independently using Claude opus 4.6, gpt-5.3-codex and gemini-3-pro-preview with the compiler oracle active, yielding 2,992 valid variants after filtering. Claude Sonnet 4.6 was used as the model for benchmark evaluation of the compiler oracle’s ability to enforce structural correctness. Each variant was scored by an automated evaluation framework along three orthogonal axes (0-100): experiment match, which measures fidelity to the source scientific content (correct reagents, concentrations, temperatures, and durations); protocol validity, which measures structural correctness as enforced by the compiler gates (unit consistency, volume conservation, and scheduling feasibility); and phase completeness, which measures whether all experimentally distinct phases of the source protocol are preserved in the generated code. The overall fidelity score is the unweighted mean of these three axes.

300 published *Nature Protocols* papers were processed through the generation pipeline. For each paper, 10 BPL program variants were generated independently using Claude opus 4.6, gpt-5.3-codex and gemini-3-pro-preview as the generation model with the BPL compiler oracle active. An automated evaluation framework scored each variant on experiment match, protocol validity, and phase completeness (0-100 each). Experiment match and phase completeness were assessed by a separate LLM evaluator (distinct from the generation model) comparing generated BPL code against the source paper content. Protocol validity was assessed deterministically by the BPL compiler. Structural determinism was assessed on a separate run of 300 protocols (10 variants each), measuring step-count stability (range, standard deviation, match percentage against a reference digest). Complexity analysis used step-count quartiles.

### Benchmark Evaluation methods

To quantify the effect of BPL on protocol standardization, we analysed both a variant-level evaluation set and the full corpus of source and processed markdown documents. Score-based analyses were performed on 2,992 processed outputs derived from 300 source papers, for which comparative protocol annotations were available. Structural analyses were performed on the same corpus. Each source document was associated with up to ten independently generated processed variants.

### Comparative protocol annotation

Each processed output was compared with its corresponding reference protocol using a structured model-assisted annotation prompt. Four scalar outputs were recorded on a 0-100 scale: experiment match, protocol validity, phase completeness and overall score. Experiment match quantified whether the processed output described the same experimental procedure as the reference protocol. Protocol validity quantified whether the output formed a coherent, executable protocol rather than malformed or nonsensical text. Phase completeness quantified whether the major experimental phases were retained, penalizing only the absence of entire major phases rather than minor wording, ordering or parameter differences.

The composite overall score was defined a priori as

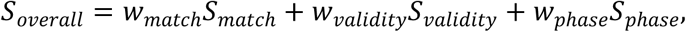

with the primary weighting

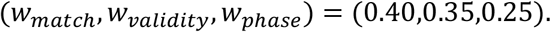

These weights were chosen to reflect task priorities rather than optimized from training data: preserving the underlying experiment was treated as the primary requirement, producing a structurally valid executable protocol was treated as the second requirement, and preserving all major protocol phases was treated as an important but slightly less fundamental requirement. Thus, experiment match received the highest weight, followed by protocol validity and then phase completeness.

For paper-level summaries, scores were averaged across all processed variants associated with the same source paper. For paper *i*, metric *m*, and *n*_*i*_ available variants, the paper-level mean was

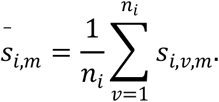

Inter-variant semantic variability was quantified as the sample standard deviation of the overall score across variants:

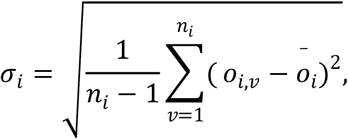

where *o*_*i,v*_ denotes the overall score of variant *v* for paper *i*. These model-derived scores were used as comparative annotations of protocol fidelity and validity rather than as an absolute ground truth measurement.

### Sensitivity analysis of score weighting

To assess whether the corpus-level conclusions depended strongly on the primary weighting scheme, we repeated the score aggregation under a grid of 47 alternative weight triplets sampled in 0.05 increments, subject to the constraints

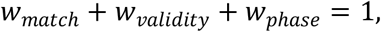

with *w*_*match*_, *w*_*validity*_ ∈ [0.20,0.60] and *w*_*phase*_ ∈ [0.15,0.45]. Across this grid, the mean variant-level overall score ranged from 93.9 to 96.6, the median score ranged from 96.0 to 97.3, and the fraction of processed variants with overall score at least 80 ranged from 97.4% to 98.5%. The mean per-paper standard deviation of the overall score ranged from 2.85 to 3.71. The alternative weighted scores remained highly correlated with the primary overall score (Pearson *r* = 0.986 to 0.999 across tested weightings). These analyses indicate that the main conclusions were insensitive to modest perturbations of the score weights.

### Inter-variant structural consistency

To quantify structural consistency across processed outputs from the same source paper, we computed the number of detected procedural steps in each processed variant and summarized the distribution of step counts within each paper. For paper *i*, the step-count range was defined as

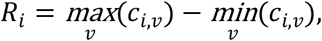

where *c*_*i,v*_ is the detected step count of variant *v*. Papers with *R*_*i*_ = 0 were classified as structurally stable, whereas papers with *R*_*i*_ > 0 were classified as structurally variable. Per-paper records from different analysis tables were harmonized by normalized paper titles before aggregation.

### Structural feature extraction

To compare natural-language source documents and processed outputs in a shared structural space, each markdown document was encoded using document-level structural features. The extracted features were word count, line count, step count, bold-format count, count of the token CRITICAL, dash-bullet count, markdown header count and the number of canonical protocol marker classes present. Step count was detected by regular expressions covering the procedural formats observed in the corpus, including heading-style steps, emphasized step markers and numbered procedural lines. Canonical protocol marker classes were defined as the presence of objective, materials, expected results, troubleshooting, critical-step language and explicit step enumeration.

### Dimensionality reduction of document structure

Principal component analysis (PCA) was used to embed documents in a low-dimensional structural feature space. For PCA, we used six count-based features: word count, line count, step count, bold-format count, CRITICAL-token count and dash-bullet count. To reduce the effect of long-tailed count distributions, each feature *x* was transformed as

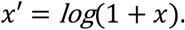

The resulting feature matrix was standardized to zero mean and unit variance before PCA, and the first two principal components were retained for visualization and distance-based summaries. For visualization only, processed outputs were additionally grouped by k-means clustering (*k* = 4) in PCA space to reveal substructure within the processed manifold; these cluster assignments were not used in the quantitative indices described below.

### Indices of structural protocolization

We summarized BPL-induced structural normalization using descriptive indices. Explicit steps (%) was defined as the proportion of documents containing at least one detected procedural step:

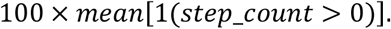

Schema coverage (%) was defined as the proportion of documents containing at least three canonical protocol marker classes:

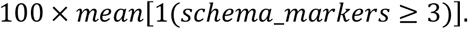

Protocol proximity was defined relative to the centroid of processed-protocol space in PCA coordinates. Let *μ*_*proc*_ denote the centroid of all processed outputs and let *d*(*p, μ*_*proc*_) = ∥ *p* − *μ*_*proc*_ ∥_2_ be the Euclidean distance between document *p* and that centroid. We defined the protocol-proximity index as

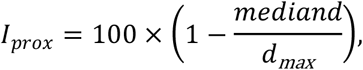

where *d*_*max*_ is the maximum observed document-to-*μ*_*proc*_ distance across the compared document classes. Higher values indicate that a document set lies closer to the processed-protocol manifold.

Structural compactness was defined from within-class concentration in PCA space. For a document class with centroid *μ*_*class*_, we computed the Euclidean distance of each document to its class centroid and summarized compactness as

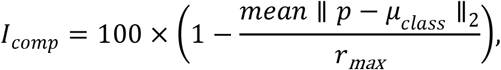

where *r*_*max*_ is the maximum observed within-class spread across the compared classes. Higher values therefore indicate a tighter, more homogeneous structural distribution. Protocol proximity and structural compactness are normalized corpus-level indices and should be interpreted as relative measures within the analysed dataset rather than as absolute percentages.

### Statistical reporting

All reported quantities are descriptive summaries of the analysed corpus. Means, medians, quartiles, standard deviations and Euclidean distances in PCA space were computed directly from the full dataset after version filtering. No formal null-hypothesis testing was applied in these analyses. All computations and visualizations were implemented in Python using NumPy, scikit-learn and Matplotlib.

### Strains and Plasmids

#### Dasher GFP Promoter Library Construction and Transformation

*E. coli* DH5α and C43(DE3) were used as the cloning and expression hosts, respectively. Eleven Dasher GFP expression plasmids, each driven by a distinct TrcT7 hybrid promoter variant (TrcT7, Trcp1 through Trcp10), were assembled by two-fragment Gibson Assembly. The backbone (~5,152 bp) was amplified using a shared forward primer (EV_F_shared, supplementary table 2) paired with 11 promoter-specific reverse primers (EV_R_PRO primers, supplementary table 2) that introduced the reverse complement of the 34-bp promoter sequence at the fragment’s 3′ end. The insert (~816 bp, encompassing the lacO operator, RBS, and Dasher GFP coding sequence, supplementary table 2) was amplified using 11 promoter-specific forward primers (insF_PRO primers, supplementary table 2) carrying the 34-bp promoter as a 5′ tail, paired with a shared reverse primer (ins_R_shared, supplementary table 2); the promoter sequence thus provided the Gibson overlap between the two fragments. PCR was performed using 2× Phanta Max Master Mix (Vazyme Biotech Co., Ltd) (98 °C 30 s; 30 cycles of 98 °C 10 s, 65 °C 20 s, 72 °C 2.5 min; 72 °C 5 min). Residual template was removed by DpnI (New England Biolabs) digestion (37 °C, 1 h), and PCR products were purified by magnetic beads (Vazyme Biotech Co., Ltd) and eluted in nuclease-free water. Gibson Assembly reactions (50 °C, 15 min) used ClonExpress Ultra One Step Cloning Kit (Vazyme Biotech Co., Ltd). All 11 constructs were prepared in parallel by a liquid handling workstation (Biomek i7) and conventional manual pipetting under identical reagent conditions, to benchmark assembly performance. Assembly products were transformed into chemically competent *E. coli* DH5α by heat shock, plated on LB agar with 50 µg mL^−1^ kanamycin, and sequence-verified by Sanger sequencing. Verified plasmids were then transformed into *E. coli* C43(DE3); transformants were plated on Q-tray LB–kanamycin plates via serial dilution spotting, and four colonies per construct were picked by an automated colony picker (QPix 420) into 96-deep-well plates containing TB medium supplemented with 50 µg mL^−1^ kanamycin for overnight seed culture.

#### GFP Expression and Fluorescence Measurement

For expression assays, 10 μL of overnight seed culture was inoculated into 390 μL TB medium supplemented with 50 μg mL^−1^ kanamycin and 10 g L^−1^ glucose in 96-deep-well plates, sealed with gas-permeable film, and incubated at 30 °C, 800 rpm. When OD_600_ reached ~0.8, expression was induced by addition of IPTG to a final concentration of 1 mM; non-induced controls received an equal volume of nuclease-free water. Cultures were incubated for a further 6 h under the same conditions. All 11 promoter constructs were assayed in four biological replicates (independent clones), with separate plates for liquid handler assisted and manual assembly groups and for induced and uninduced conditions (four plates total). After induction, 20 μL of each culture was diluted 10-fold in nuclease-free water and transferred to a clear-bottom 96-well microplate. OD_600_ and GFP fluorescence (Ex 485/20 nm, Em 528/20 nm, gain 35, bottom optics) were measured on a multi-mode microplate reader (Biotek Synergy S1LFA HTX Multi-Mode Reader). Fluorescence signals were normalized using four biological replicates for each sample, without calculating the RFU/OD_600_ ratio. Both OD_600_ and fluorescence values were individually normalized based on their respective four replicates.

#### Chromatographic method translation and validation

To validate the method translation, standard solutions of retinol, retinal, retinyl acetate, lycopene, and β-carotene (50 mg/L) were prepared and analyzed on an Agilent UHPLC 1290 Infinity II equipped with diode-array detection. The reference HPLC method utilized an Agilent ValueLab LC C18 column (170 Å, 4.6 × 250 mm, 5 μm) running a 32-minute gradient of water and acetonitrile/isopropanol (75:25, v/v) at 1.0 mL/min with a 10 μL injection volume. BPL-COGEN translated this protocol for a Phenomenex Kinetex C18 core-shell column (100 Å, 2.1 × 50 mm, 1.7 μm) by applying scaling formulae from the 2025 Chinese Pharmacopoeia General Chapter 0512 (Chinese Pharmacopoeia, 2025). Guided by the compiler’s calculated parameters and compliance warnings, the executed UHPLC method adopted a 0.6 mL/min flow rate and a 1 μL injection volume.

## Data availability

The Nature protocol Benchmark are available on HuggingFace Hub at https://huggingface.co/datasets/BotaBio/BPL-EVAL

## Code availability

All software is available at https://gitlab.com/bota-biosciences-public/bpl-code under MIT license, including the compiler (bplc), BPL-COGEN pipeline, corpus generation pipeline, and fine-tuning scripts.

## Notes

### Competing Interest Statement

The authors have declared no competing interest.

