## Supplementary note for "Towards autonomous biology: Compiler-Verified Protocols as a Foundation for Real-World AI Execution"

#### Biology protocol language

**BPL Program structure.** A BPL program consists of a declaration section followed by one or more protocol blocks. Declarations define the material world — reagents with physical properties, stocks with concentrations, labware with capacity constraints and execution targets (human or robotic). The protocol block contains numbered steps, each mapping to a single laboratory intent. The grammar (463 lines, Lark PEG specification) enforces this structure:

```
reagent Tryptone  { physical_form: Solid, MW: 145.19 g/mol, cas: "91079-40-2" }
reagent NaCl      { physical_form: Solid, MW: 58.44 g/mol }
reagent ROWater   { physical_form: Liquid }


stock LB_Base {
  reagent: Tryptone
  concentration: 10 g/L
  solvent: ROWater
}


labware Beaker1L      { footprint: Custom, well_volume: 1200 mL }
labware GlassBottle1L { footprint: Custom, well_volume: 1000 mL }


@target(human)
@expected_time(2 h)
protocol PrepareLBMedia(final_volume: Volume = 1000 mL) {
  @step(1, "Add RO water to beaker")
  add(ROWater, volume: 800 mL, to: Beaker1L)

  @step(2, "Weigh and add tryptone and NaCl")
  parallel {
    add(Tryptone, mass: 10 g, to: Beaker1L)
    add(NaCl, mass: 10 g, to: Beaker1L)
```

```
  }

  @step(3, "Mix until fully dissolved")
  mix(target: Beaker1L, cycles: 10, method: stir)

  @step(4, "Adjust to final volume")
  transfer(from: ROWater, to: Beaker1L, volume: 200 mL)

  @step(5, "Transfer to autoclave bottle")
  transfer(from: Beaker1L, volume: final_volume, to: GlassBottle1L)

  @step(6, "Autoclave at 121 °C for 20 min")
  incubate(target: GlassBottle1L, temperature: 121 degC, time: 20 min)
}
```

**Biology-native type system and dimensional analysis.** BPL implements compile-time dimensional analysis across nine base dimensions: volume, mass, temperature, time, concentration, amount (moles), pressure, speed and length. Every quantity literal carries a mandatory unit annotation; the compiler tracks dimensions through all arithmetic expressions and rejects operations on incompatible types at compile time:

```
transfer(from: source, to: dest, volume: 100 uL)   # valid: volume dimension
add(NaCl, mass: 10 g, to: Beaker1L)                # valid: mass dimension
incubate(target: plate, temperature: 37 degC, time: 1 h)  # valid: temperature, time

# Compile-time errors:
# transfer(from: A, to: B, volume: 100 uL + 50 mg)  → dimension mismatch
# incubate(target: X, temperature: 37 mL)           → expected temperature, got volume
```

Supported units span the laboratory domain: mL/uL/nL (volume), g/mg/ug/ng (mass), degC (temperature), M/mM/uM (molar concentration), g/L/mg/mL (mass concentration), min/s/h (time), rpm/rcf (centrifugation speed) and fold factors (100X, 1000X). Compound dimensions are derived automatically — concentration as mass/volume, rate as volume/time — and enforced through the expression type checker.

**Laboratory intents and multi-target lowering.** BPL defines ~14 high-level laboratory intents, each representing a conceptual operation rather than a machine instruction:

| Category | Intents |
|---|---|
| Liquid handling | transfer, add, distribute, consolidate |
| Physical processes | mix, incubate, centrifuge |
| Specialized | run_pcr, pick_colonies, adjust_ph, heat_shock, on_ice |
| Manual / control | manual (with StateDelta), checkpoint |

Each intent accepts typed named arguments (e.g., transfer(from: <container>, to: <container>, volume: <Volume>)). The compiler lowers intents to ~20 execution primitives (ASPIRATE, DISPENSE, PICK_TIPS, DROP_TIPS, SET_TEMPERATURE, READ_ABSORBANCE, MOVE_LABWARE, etc.), enabling the same BPL source to compile to human-readable worklists, robotic control sequences or virtual simulations. This intent–primitive separation means that a single protocol definition adapts to different execution backends without source modification.

**Container state tracking and capacity enforcement.** Every container in a BPL program is a stateful entity tracked by the compiler from declaration through every operation. Containers are referenced symbolically (source, dest, pcr_plate:A1) with compile-time tracking of current volume, concentration and maximum capacity. When a transfer executes, the compiler automatically decrements the source volume and increments the destination volume; if the destination would exceed its declared capacity, or the source would go negative, the program is rejected at compile time:

```
labware SmallTube { footprint: Tube, well_volume: 500 uL }


# The following would produce a compile-time capacity error:

# transfer(from: reservoir, to: SmallTube, volume: 800 uL)

# → Error: transfer of 800 uL exceeds SmallTube capacity (500 uL)
```

For operations not automatically tracked (e.g., a manual centrifugation step that removes supernatant), StateDelta expressions declare the state effects explicitly, maintaining the compiler's volume accounting across all steps.

**Trust model and compliance.** A three-tier trust model (Declared → Calibrated → Verified) tracks the provenance and confidence of every state value. *Declared* values are user- or compiler-asserted; *Calibrated* values are confirmed by instrument measurement (balance, pipette, plate reader); *Verified* values carry an operator signature with a hash-chained audit trail, satisfying GLP/GMP requirements for 21 CFR Part 11 compliance.

**Control flow.** BPL supports if/else conditional branching, for/while loops and parallel { } blocks for concurrent operations. Recovery handlers (@on_error) and invariant blocks provide structured error handling tied to specific failure modes.

### Three-phase BPL language design process

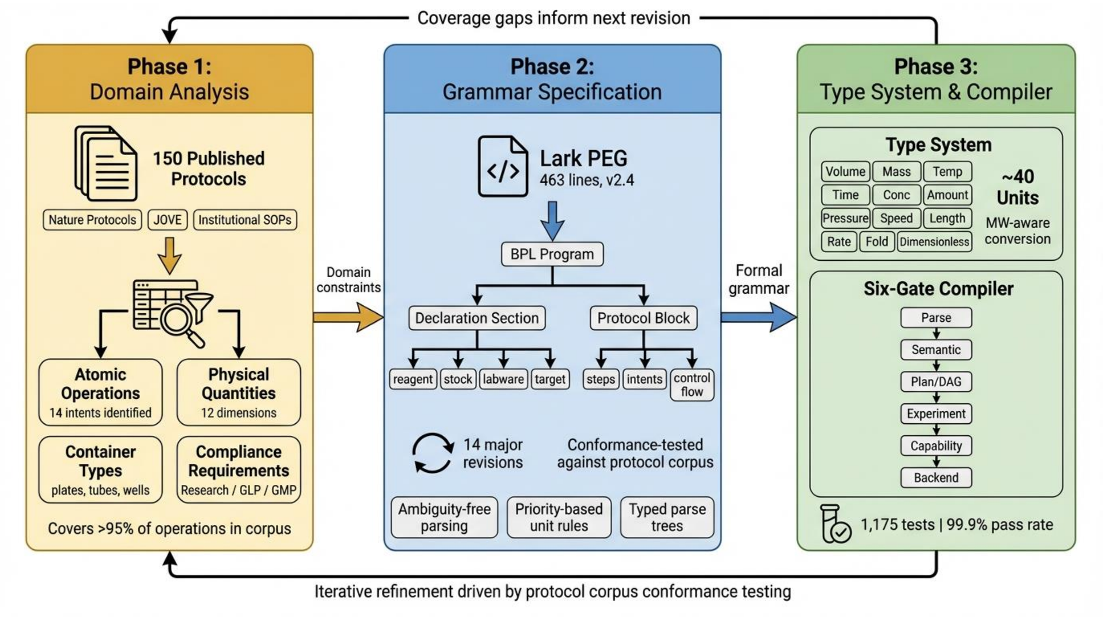


**Fig. S1. Three-phase BPL language design process.** Phase 1 (Domain Analysis): 150 published protocols from *Nature Protocols*, JOVE, and institutional repositories were systematically analyzed to identify 14 atomic laboratory intents, 12 physical dimensions, container types, and compliance requirements covering >95% of operations in the corpus. Phase 2 (Grammar Specification): Domain constraints were encoded as a 463-line Lark PEG grammar (v2.4), defining a two-section program structure (declarations and protocol blocks) refined through 14 major revisions driven by conformance testing. Phase 3 (Type System and Compiler): A biology-native type system with ~40 units and molecular-weight-aware concentration conversion was implemented alongside a six-gate compiler pipeline validated by 1,175 tests (99.9% pass rate). A feedback loop feeds coverage gaps back from compilation to domain analysis, driving iterative refinement.

### Four-layer quality assurance pipeline for training corpus generation

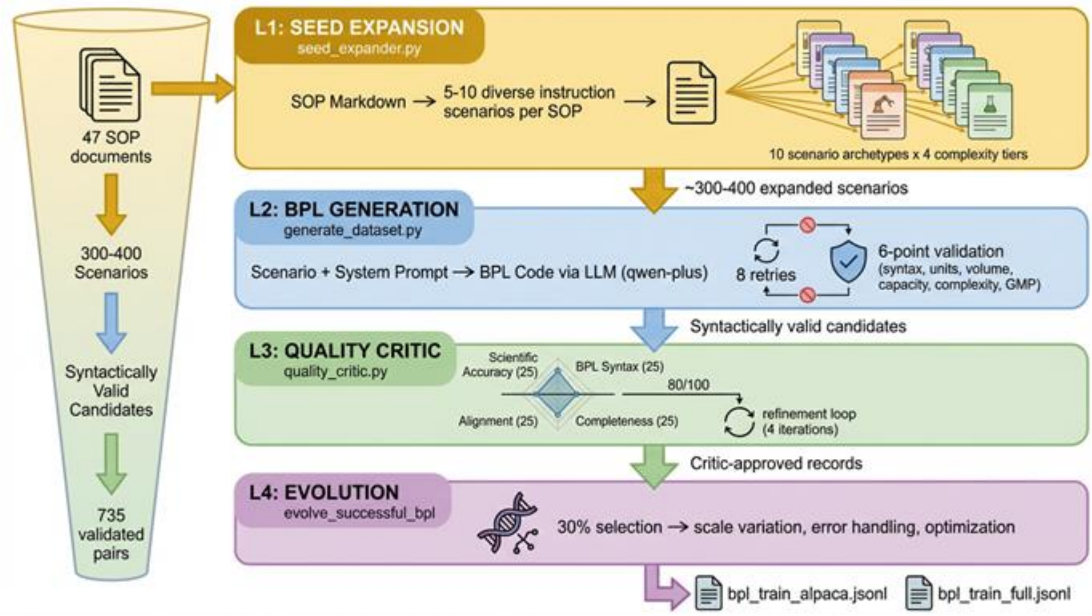


**Fig. S2. Four-layer quality assurance pipeline for training corpus generation**. Seed expansion generates diverse instruction scenarios from 47 SOPs. BPL generation produces candidate code with 6-point compiler validation. The LLM quality critic scores candidates on four axes (threshold 80/100) and routes failures through a refinement loop. Evolutionary variation diversifies 30% of approved records. Compound filtering yields 2,714 validated pairs (~24%).

### Diagnostics and repair examples

**Supplementary Table 1. Compiler diagnostic categories, repair hints, and repaired examples.**

| Category | Example (fails) | Compiler repair hint | Repaired example (passes) |
|---|---|---|---|
| Dimension mismatch | transfer(from: NaCl, to: Tube, volume: 50 mg) | "Expected volume dimension; got mass. Use mL, uL, or nL." | transfer(from: NaCl, to: Tube, volume: 50 uL) |
| Capacity violation | transfer(from: reservoir, to: Small Tube, volume: 800 uL) where Small Tube capacity = 500 uL | "Transfer of 800 uL exceeds Small Tube capacity (500 uL). Split transfer or use larger container." | transfer(from: reservoir, to: Large Tube, volume: 800 uL) with labware Large Tube { well volume: 1500 uL } |
| Undeclared identifier | add(Tryptone, mass: 10 g, to: Beaker) with no reagent Tryptone declaration | "Identifier 'Tryptone' not found in declarations. Add a reagent declaration." | Add reagent Tryptone { physical form: Solid, MW: 145.19 g/mol } to declarations |
| State conflict | transfer(from: Source, to: Dest, volume: 200 uL) when Source has 150 uL remaining | "Transfer would leave source at -50 uL. Check transfer volume or add replenishment step." | transfer(from: Source, to: Dest, volume: 150 uL) or add prior add(ROWater, volume: 100 uL, to: Source) |
| Trust violation | incubate(target: Plate, temperature: 37 degC) in @compliance(GMP) context with Declared trust | "GMP context requires Verified trust level; current trust is Declared. Add calibration step." | Add calibrate(target: Plate, property: temperature, instrument: TempProbe) before incubation |

### Supplementary Table 2. DNA sequences used in this study

| Name | Type | Sequences |
|---|---|---|
| LacO_RBS_DasherGFP | Gene | TAGGGGAATTGTGAGCGGATAACAATTCCCCTCTAGAAATAAT<br>TTTGTTTAACTTTAAGAAGGAGATATACCatgaccgcactaacagaag<br>gagctaaactattcgaaaaggagattccttacattacagaattagagggtgatgtcgaag<br>gaatgaaattcattatcaagggcgagggtactggtgacgctactaccggtacgattaaag<br>caaagtacatctgtacaacaggtgaccttcctgttccgtgggctactctggtgagcactttgt<br>cttatggagttcaatgttttgctaaatacccttcgcacattaaagactttttcaaaagtgcaat<br>gcctgagggctatactcaggagagaacaatatctttcgaaggagatggtgtgtataaga<br>ctagggctatggtcacgtatgaaagaggatccatctacaatagagtaactttaactggtga<br>aaacttcaaaaaggacggtcacatccttagaaagaatgttgcctttcaatgcccaccatc<br>catcttgtacattttgccagacacagttaacaatggtatcagagttgagtttaaccaagctta<br>tgacatagagggtgtcaccgaaaagttggttacaaaatgttcacagatgaatcgtcccct<br>ggcaggatcagctgccgtccatatcccacgttaccatcatatcacttatcataccaagctgt<br>ccaaagatcgtgatgagagaagggatcacatgtgtttggttgaagtggtaaaggccgtg<br>gatttggatacttaccaaggttga |
| TrcT7 | Promoter | TTGACAATTAATCATCCGGCTCGTATAATGTAGG |
| Trcp1 | Promoter | TTGACAATTAATCATCCGGCTCGTATTATGTAGG |
| Trcp2 | Promoter | TTGACAATTAATCATCCGGCTCGTACTATGTAGG |
| Trcp3 | Promoter | TTGACGATTAATCATCCGGCTCGTATAATGTAGG |
| Trcp4 | Promoter | TTGACAATTAATCATCCGGCTCGTACAATGTAGG |
| Trcp5 | Promoter | TTGACAATTAATCATCCGGCTCGGATAATGTAGG |
| Trcp6 | Promoter | TTGACAATTAATCATCCGGCTCGGACAATGTAGG |
| Trcp7 | Promoter | TTTACAATTAATCATCCGGCTCGTATTATGTAGG |
| Trcp8 | Promoter | TTGATAATTAATCATCCGGCTCGTATTATGTAGG |
| Trcp9 | Promoter | TTTACGATTAATCATCCGGCTCGTATAATGTAGG |
| Trcp10 | Promoter | CTTATAATTAATCATCCGGCTCGGATTATGTAGG |
| EV_F_shared | Primer | CTCGAGCACCACCACCAC |
| ins_R_shared | Primer | AGTGGTGGTGGTGGTGGTGCTCGAGTCAACCTTGGTAAGTA<br>TCCAAATCC |
| insF_PRO321_TrcT7 | Primer | TTGACAATTAATCATCCGGCTCGTATAATGTAGGGGAATTGT<br>GAGCGGATAACAAT |
| insF_PRO330_Trcp1 | Primer | TTGACAATTAATCATCCGGCTCGTATTATGTAGGGGAATTGT<br>GAGCGGATAACAAT |
| insF_PRO331_Trcp2 | Primer | TTGACAATTAATCATCCGGCTCGTACTATGTAGGGGAATTGT<br>GAGCGGATAACAAT |
| insF_PRO332_Trcp3 | Primer | TTGACGATTAATCATCCGGCTCGTATAATGTAGGGGAATTGT<br>GAGCGGATAACAAT |
| insF_PRO333_Trcp4 | Primer | TTGACAATTAATCATCCGGCTCGTACAATGTAGGGGAATTGT<br>GAGCGGATAACAAT |
| insF_PRO334_Trcp5 | Primer | TTGACAATTAATCATCCGGCTCGGATAATGTAGGGGAATTGT<br>GAGCGGATAACAAT |
| insF_PRO335_Trcp6 | Primer | TTGACAATTAATCATCCGGCTCGGACAATGTAGGGGAATTG<br>TGAGCGGATAACAAT |
| insF_PRO336_Trcp7 | Primer | TTTACAATTAATCATCCGGCTCGTATTATGTAGGGGAATTGT<br>GAGCGGATAACAAT |
| insF_PRO337_Trcp8 | Primer | TTGATAATTAATCATCCGGCTCGTATTATGTAGGGGAATTGT<br>GAGCGGATAACAAT |

| | | |
|---|---|---|
| insF_PRO338_Trcp9 | Primer | TTTACGATTAATCATCCGGCTCGTATAATGTAGGGGAATTGT GAGCGGATAACAAT |
| insF_PRO339_Trcp10 | Primer | CTTATAATTAATCATCCGGCTCGGATTATGTAGGGGAATTGT GAGCGGATAACAAT |
| EV_R_PRO321 | Primer | CCTACATTATACGAGCCGGATGATTAATTGTCAACCTCTACG CCGGACGCAT |
| EV_R_PRO330 | Primer | CCTACATAATACGAGCCGGATGATTAATTGTCAACCTCTACG CCGGACGCAT |
| EV_R_PRO331 | Primer | CCTACATAGTACGAGCCGGATGATTAATTGTCAACCTCTACG CCGGACGCAT |
| EV_R_PRO332 | Primer | CCTACATTATACGAGCCGGATGATTAATCGTCAACCTCTACG CCGGACGCAT |
| EV_R_PRO333 | Primer | CCTACATTGTACGAGCCGGATGATTAATTGTCAACCTCTACG CCGGACGCAT |
| EV_R_PRO334 | Primer | CCTACATTATCCGAGCCGGATGATTAATTGTCAACCTCTACG CCGGACGCAT |
| EV_R_PRO335 | Primer | CCTACATTGTCCGAGCCGGATGATTAATTGTCAACCTCTACG CCGGACGCAT |
| EV_R_PRO336 | Primer | CCTACATAATACGAGCCGGATGATTAATTGTAAACCTCTACG CCGGACGCAT |
| EV_R_PRO337 | Primer | CCTACATAATACGAGCCGGATGATTAATTATCAACCTCTACG CCGGACGCAT |
| EV_R_PRO338 | Primer | CCTACATTATACGAGCCGGATGATTAATCGTAAACCTCTACG CCGGACGCAT |
| EV_R_PRO339 | Primer | CCTACATAATCCGAGCCGGATGATTAATTATAAGCCTCTACG CCGGACGCAT |
