## Supplementary file 9 for "Towards autonomous biology: Compiler-Verified Protocols as a Foundation for Real-World AI Execution"

### Supplementary Information: BPL-COGEN HPLC-to-UHPLC Method Translation

This file provides the generated BPL program, deterministic execution plan, and full diagnostic report referenced in the manuscript.

The reported values are the compiler-generated method-translation values. As described in the Methods, the bench execution used practical instrument rounding: 0.613 mL/min to 0.6 mL/min, 0.420 µL to 1 µL, and 2.18 min to approximately 2.1 min.

#### Supplementary File Inventory

| File | Contents |
|---|---|
| hplc_to_uhplc_bpl_program.bpl | Parser-valid BPL source listing for the translated method |
| hplc_to_uhplc_execution_plan.json | Machine-readable deterministic execution plan |
| hplc_to_uhplc_diagnostic_report.json | Machine-readable full diagnostic report |

#### Source and Target Methods

| Field | Source HPLC | Target UHPLC |
|---|---|---|
| Column | Agilent ValueLab LC C18 170A | Phenomenex Kinetex C18 100A |
| Dimensions | 4.6 × 250 mm | 2.1 × 50 mm |
| Particle size | 5.0 µm | 1.7 µm |
| Flow rate | 1.000 mL/min | 0.613 mL/min |
| Injection volume | 10.000 µL | 0.420 µL |
| Run time | 32.00 min | 2.18 min |
| Solvent volume | 32.000 mL | 1.336 mL |
| Speed factor | - | 14.679× |
| Solvent saving | - | 95.824% |

#### Translated Gradient

| Time (min) | %A | %B |
|---|---|---|
| 0.00 | 25.0 | 75.0 |
| 0.37 | 25.0 | 75.0 |
| 0.51 | 0.0 | 100.0 |
| 2.04 | 0.0 | 100.0 |
| 2.18 | 25.0 | 75.0 |

#### Supplementary Program S1. Generated BPL Program

```
// Supplementary BPL Program S1
// BPL-COGEN generated HPLC-to-UHPLC method translation for Fig. 5.
// The chromatography conversion engine applies ChP 0512 column-scaling rules.

intent define_column(id: String, name: String, inner_diameter: Number, length: Number, particle_size: Number, pore_size_angstrom: Number, packing: String, max_pressure: Number)
intent define_method(id: String, column: String, flow_rate: Number, injection_volume: Number, temperature: Number, detection_mode: String, wavelengths_nm: List<Number>, mobile_phase_a: String, mobile_phase_b: String)
intent gradient_point(method: String, time: Number, percent_b: Number)
intent translate_method(source_method: String, target_column: String, standard: String, flow_rate: Number, injection_volume: Number, run_time: Number, speed_factor: Number, solvent_saving_percent: Number, estimated_pressure_ratio: Number)
intent record_diagnostic(parameter: String, severity: String, message: String)
intent require_verification(check: String)

@target(human)
@expected_time(2.18 min)
protocol TranslateHPLCToUHPLCMethod {
    define_column(id: "valuelab-c18-4.6x250-5um", name: "Agilent ValueLab LC C18 170A", inner_diameter: 4.6 mm, length: 250 mm, particle_size: 5.0 um, pore_size_angstrom: 170, packing: "fully_porous", max_pressure: 400 bar)

    define_method(id: "source_hplc", column: "valuelab-c18-4.6x250-5um", flow_rate: 1.000 mL/min, injection_volume: 10.000 uL, temperature: 50.0 degC, detection_mode: "DAD", wavelengths_nm: [328, 382, 452, 475, 210], mobile_phase_a: "Ultrapure water", mobile_phase_b: "Acetonitrile/Isopropanol 75:25")

    define_column(id: "kinetex-c18-2.1x50-1.7um", name: "Phenomenex Kinetex C18 100A", inner_diameter: 2.1 mm, length: 50 mm, particle_size: 1.7 um, pore_size_angstrom: 100, packing: "superficially_porous", max_pressure: 1000 bar)

    translate_method(source_method: "source_hplc", target_column: "kinetex-c18-2.1x50-1.7um", standard: "Chinese Pharmacopoeia 2025 General Chapter 0512", flow_rate: 0.613 mL/min, injection_volume: 0.420 uL, run_time: 2.18 min, speed_factor: 14.679, solvent_saving_percent: 95.824 percent, estimated_pressure_ratio: 5.089)

    define_method(id: "translated_uhplc", column: "kinetex-c18-2.1x50-1.7um", flow_rate: 0.613 mL/min, injection_volume: 0.420 uL, temperature: 50.0 degC, detection_mode: "DAD", wavelengths_nm: [328, 382, 452, 475, 210], mobile_phase_a: "Ultrapure water", mobile_phase_b: "Acetonitrile/Isopropanol 75:25")

    gradient_point(method: translated_uhplc, time: 0.00 min, percent_b: 75.0 percent)
    gradient_point(method: translated_uhplc, time: 0.37 min, percent_b: 75.0 percent)
    gradient_point(method: translated_uhplc, time: 0.51 min, percent_b: 100.0 percent)
    gradient_point(method: translated_uhplc, time: 2.04 min, percent_b: 100.0 percent)
    gradient_point(method: translated_uhplc, time: 2.18 min, percent_b: 75.0 percent)

    record_diagnostic(parameter: "L/dp", severity: "warning", message: "Target L/dp (29412) is 41% below source (50000). ChP 0512 allows max -25%.")
    record_diagnostic(parameter: "packing_type", severity: "warning", message: "Packing change: fully_porous -> superficially_porous. Per ChP 0512, selectivity and elution order must remain unchanged.")
    record_diagnostic(parameter: "pore_size", severity: "warning", message: "Pore category change: peptide -> small_molecule. May affect retention and mass transfer for large analytes.")
    record_diagnostic(parameter: "backpressure", severity: "warning", message: "Estimated pressure increase: 5.1x. Target column rated to 1000 bar. Verify instrument pressure capability.")
    record_diagnostic(parameter: "column_temp", severity: "info", message: "Temperature: 50.0C (adjustable +/-5.0C per ChP gradient rules)")
    record_diagnostic(parameter: "detection_wavelength", severity: "info", message: "Detection wavelengths must not change per ChP 0512.")

    require_verification(check: "Verify elution order and resolution on target UHPLC column before routine use.")
    require_verification(check: "Verify instrument pressure headroom for estimated 5.1x backpressure increase.")
}
```

#### Supplementary Data S1. Execution Plan

| Step | Name | Primitive | Dependencies |
|---|---|---|---|
| step_0001 | normalize_source_method | NORMALIZE_METHOD | - |
| step_0002 | resolve_target_column | RESOLVE_COLUMN | step_0001 |
| step_0003 | check_chp0512_column_constraints | COMPLIANCE_CHECK | step_0002 |
| step_0004 | scale_flow_rate | CALCULATE | step_0003 |
| step_0005 | scale_injection_volume | CALCULATE | step_0003 |
| step_0006 | scale_gradient_timetable | CALCULATE_GRADIENT | step_0004, step_0005 |
| step_0007 | estimate_backpressure | PRESSURE_ESTIMATE | step_0004 |
| step_0008 | emit_verified_method_package | EMIT_ARTIFACTS | step_0006, step_0007 |

#### Supplementary Data S2. Full Diagnostic Report

Validation gates

| Gate | Status | Diagnostics |
|---|---|---|
| parse | passed | 0 warning, 0 info |
| semantic | passed | 0 warning, 0 info |
| method_conversion | passed_with_warnings | 4 warning, 2 info |
| artifact_emission | passed | 0 warning, 0 info |

Calculation audit

| Check | Result |
|---|---|
| Flow scaling | F2 = F1 * (dc2^2 * dp1) / (dc1^2 * dp2); result = 0.613 mL/min |
| Injection scaling | Vinj2 = Vinj1 * (L2 * dc2^2) / (L1 * dc1^2); result = 0.420 µL |
| Pressure estimate | 5.089× target/source |
| Solvent use | 32.000 mL → 1.336 mL; saving = 95.824% |

Diagnostic listing

| Severity | Parameter | Diagnostic |
|---|---|---|
| warning | L/dp | Target L/dp (29412) is 41% below source (50000). ChP 0512 allows max -25%. |
| warning | packing_type | Packing change: fully_porous -> superficially_porous. Per ChP 0512, selectivity and elution order must remain unchanged. |
| warning | pore_size | Pore category change: peptide -> small_molecule. May affect retention and mass transfer for large analytes. |
| warning | backpressure | Estimated pressure increase: 5.1x. Target column rated to 1000 bar. Verify instrument pressure capability. |
| info | column_temp | Temperature: 50.0C (adjustable +/-5.0C per ChP gradient rules) |
| info | detection_wavelength | Detection wavelengths must not change per ChP 0512. |
