## Supplementary file 2 for "Towards autonomous biology: Compiler-Verified Protocols as a Foundation for Real-World AI Execution"

**GFP_Promoter_Library_Gibson_Assembly_—Automation_protocols**

**1. Experiment Overview**

This experiment aims to construct a DasherGFP expression library driven by 11 different promoters. The study focuses on using the single shared template DNA3776 to amplify both the vector backbone (Backbone) and the target insert fragment (Insert) containing different promoter variants. The assembly task combines high-fidelity PCR with Gibson Assembly, automated on the Beckman Biomek I7. Successful completion of this assembly will establish the foundation for downstream fluorescence expression screening and quantification.

**2. Experimental Design**

To address the above assembly requirements, we designed a 2-fragment Gibson Assembly strategy. [ASSUMPTION: Template DNA3776 purity reaches sequencing grade and contains no PCR inhibitors, suitable for direct use as the base PCR template]. Primers have been previously designed and verified for physical parameters, ensuring no significant secondary structures or hairpins. The promoter sequences (34 bp) serve as the continuous Gibson Assembly overlap region, directly introduced into the amplification product termini.

**2.1 Construct Association Matrix**

Based on vector pBEE (EV2442) and DasherGFP (DNA14963), the following 11 target constructs will be generated:

| **Target Plasmid Clone** | **Target Promoter** | **Vector Backbone** |
| --- | --- | --- |
| pBEE_TrcT7_DasherGFP | PRO321 | EV2442 |
| pBEE_Trcp1_DasherGFP | PRO330 | EV2442 |
| pBEE_Trcp2_DasherGFP | PRO331 | EV2442 |
| pBEE_Trcp3_DasherGFP | PRO332 | EV2442 |
| pBEE_Trcp4_DasherGFP | PRO333 | EV2442 |
| pBEE_Trcp5_DasherGFP | PRO334 | EV2442 |
| pBEE_Trcp6_DasherGFP | PRO335 | EV2442 |
| pBEE_Trcp7_DasherGFP | PRO336 | EV2442 |
| pBEE_Trcp8_DasherGFP | PRO337 | EV2442 |
| pBEE_Trcp9_DasherGFP | PRO338 | EV2442 |
| pBEE_Trcp10_DasherGFP | PRO339 | EV2442 |

**2.2 PCR Amplification Strategy**

- **Backbone PCR (5,152 bp)**: Uses universal forward primer EV_F_shared and promoter-specific reverse primer EV_R_PROxxx. The target fragment will contain the reverse complement region of the promoter (RC Promoter 34 bp).
- **Insert PCR (816 bp)**: Uses promoter-specific forward primer insF_PROxxx and universal reverse primer ins_R_shared. The specific primer contains the complete promoter region and the downstream lacO sequence (22 bp).

**3. Data Collection and QC**

All core products will generate auditable records at process checkpoints. Operators will perform 1% agarose gel electrophoresis on all PCR reactions. Backbone target band is expected at ~5.1 kb; Insert expected at ~800 bp. After transformation and colony isolation, ≥3 clones per group will be randomly selected for colony PCR, with an expected target band of 816 bp. Sanger or whole-plasmid nanopore sequencing will cover assembly junctions, promoter, and coding regions to confirm multi-sequence integrity.

**4. Statistical Analysis Plan**

Given the specific nature of this gene fragment assembly process, randomized controls and blinding interventions are not applicable (marked N/A).

- **Primary Comparison Metric**: Sequence match percentage between picked recombinant clone sequences and the reference in silico vector map.
- **Missing Data Handling**: If a construct yields 0 colonies or all clones test as false positives, the process will be treated as complete loss. Strategy is to retry the original step and supplement analysis, not imputation.
- **Sensitivity Analysis**: Separate evaluation for sequencing errors vs. actual mutations.

**5. Materials and Resources**

- **Polymerase System**: Phanta Max Master Mix (Vazyme Biotech Co., Ltd). For the convenience of pipetting, the Master Mix is diluted with diH_2_O in a ratio of Mater Mix : water = 25 : 19.
- **Assembly Reagent**: Gibson Assembly 2× Master Mix, ClonExpress Ultra One Step Cloning Kit (Vazyme Biotech Co., Ltd)
- **Auxiliary Enzyme**: DpnI restriction enzyme (New England Biolabs)
- **Competent Cell System**: DH5α chemically competent E. coli, with SOC as recovery medium.
- **Consumables**: PCR purification kit (magnetic beads method, Vazyme Biotech Co., Ltd), 96-well PCR plates, and LB plates with 50 μg/mL Kanamycin.
- **Equipment**

| **Equipment** | **Notes** |
| --- | --- |
| Single-channel pipettes: P2, P10, P20, P200 |  |
| Multi-channel pipette (optional): P10, P200 | For parallel dispensing |
| Thermal cycler | Standard 96-well block |
| 42°C water bath or heat block | For heat-shock transformation |
| 37°C shaking incubator (225 rpm) | For SOC recovery |
| NanoDrop | For DNA quantification |
| Gel electrophoresis system | For PCR result validation |
| Microcentrifuge | For spin-down |
| Beckman Biomek I7 | For automated liquid handling |

**6. Success Criteria and Decision Rules**

1. **Electrophoresis Amplification Success**: Main gel band concentration ≥80%; tailing or non-specific amplification judged as secondary.
2. **Plasmid Positive Prediction**: Post-sequencing precise match rate should be 100% (no mutations outside promoter equivalency).

**7. Primer Description**

All 24 primers are standard desalt purification at 100 nmol synthesis scale (all lengths ≤60 bp). See supplementary table 2 for full sequences.

**8. 96-Well Plate Layout Design**

- **Layout 1: Primer Plate (PrimerPlate_DasherGFP)**
- Wells A1–A12: P01 through P12 (10 μM working solution)
- Wells B1–B12: P13 through P24 (10 μM working solution)
- **Layout 2: Template Plate (TemplatePlate_DasherGFP)**
- Well A1: Universal template DNA3776 stock (~1 ng/μL)
- **Layout 3: PCR Plate (PCRPlate_DasherGFP)**
- Backbone amplification Row A: A1 (PRO321), A2 (PRO330), A3 (PRO331)... A11 (PRO339)
- Insert amplification Row B: B1 (PRO321), B2 (PRO330), B3 (PRO331)... B11 (PRO339)
- **Layout 4: Assembly Plate (GibsonPlate_DasherGFP)** and **Transformation Plate (TransformPlate_DasherGFP)**
- Aligned with PCR plate positions, using only Row A wells A1–A11.

**9. Beckman Biomek Worklists**

All downstream CSV tables feed directly into the workstation transfer command execution queue.

**Worklist 1: Primer Cherrypick → PCR Plate (Primer_to_PCR_Cherrypick.csv)**

Transfer: Extract and assemble primers required for Backbone and Insert (44 total aspirations). Volume: 2.5 μL each.

*(Full 44-row CSV available in companion Worklist 1 document)*

**Worklist 2: Template Cherrypick → PCR Plate (Template_to_PCR_Cherrypick.csv)**

Distribute same template across all wells. 22 transfers, 1 μL each.

**Worklist 3: PCR Master Mix Dispense (PCR_MasterMix_Dispense.csv)**

Add buffer system and enzyme to activate amplification. From Reagent Plate A1 to all 22 PCR wells, 44 μL each.

**Worklist 4: PCR Products → Gibson Assembly Plate (PCR_to_Gibson_Cherrypick.csv)**

Combine corresponding Backbone (Row A) and Insert (Row B) (2.5 μL each) into the new assembly plate. 22 transfers → 11 Gibson wells.

**Worklist 5: Gibson Master Mix Dispense (Gibson_MasterMix_Dispense.csv)**

Add equal volume (5 μL) of Gibson Assembly 2× Master Mix. 11 transfers.

**Worklist 6: Gibson Products → Transformation Plate (Gibson_to_Transform.csv)**

Transfer 2 μL assembly product to E. coli transformation system. 11 transfers.

**10.** **Step-by-Step Experiment Protocol with Automation**

**Step 1: Primer Reconstitution**

- Centrifuge primer tubes to pellet lyophilized powder.
- Add calculated volume of nuclease-free water or TE Buffer to prepare stock (100 μM).
- Vortex 3–5 seconds.
- Transfer 10 μL to dilution tube; add 90 μL water for working solution (10 μM).
- Aliquot into designated primer plate well positions.

**Step 2: PCR Amplification**

- Thaw pre-prepared high-fidelity PCR 2× Master Mix at room temperature; place in reagent reservoir.
- Run Worklist 1: Primer Cherrypick transfer.
- Run Worklist 2: Template distribution.
- Run Worklist 3: Master mix dispensing.
- Seal plate and transfer to thermal cycler:
- 98°C initial denaturation 30 s.
- 30 cycles: 98°C denaturation 10 s → 65°C annealing 20 s → 72°C extension 2.5 min.
- 72°C final extension 5 min → Hold at 4°C.
- Run 1% agarose gel electrophoresis for QC. Load 5 μL of each PCR reaction + 1 μL 6× loading dye onto 1% agarose gel. Run at 120 V for 25–30 min.

**Step 3: PCR Product Purification**

- Mix 2 μL of DpnI (40U) to 50 μL PCR product and carry out digestion at 37°C for 1 h to eliminate template background.
- Purify DNA fragments using magnetic beads PCR cleanup kits following manufacturer’s protocol (Vazyme Biotech Co., Ltd). Elute in 50 μL nuclease-free water.
- Measure concentration with NanoDrop.
- Transfer to fresh 96-well plate for isolated storage.

**Step 4: Gibson Assembly Automated Incubation**

- Run Worklist 4: Pair backbone and insert fragments.
- Run Worklist 5: Dispense Gibson Assembly master mix.
- Remove and incubate at 50°C for 15 min in water bath or thermal block.
- Return assembly plate to ice or 4°C for rapid cooling.

**Step 5: E. coli Chemical Heat-Shock Transformation**

- Thaw DH5α cells on ice. Distribute 50 μL aliquots of DH5α competent cells on ice into well A1-A11 in a 96-well plate.
- Run Worklist 6: Transfer 2 μL Gibson product to competent cell suspension.
- Incubate on ice 30 min.
- Heat-shock at 42°C for 30 seconds.
- Return to ice for 2 min.
- Add 950 μL room temperature SOC recovery medium.
- Incubate at 37°C, 225 rpm for 1 hour.
- Spread 100 μL of the recovered cells on LB + 50 μg/mL Kanamycin plates; incubate inverted at 37°C for 14–16 h.
