## Supplementary file 1 for "Towards autonomous biology: Compiler-Verified Protocols as a Foundation for Real-World AI Execution"

**GFP_Promoter_Library_Gibson_Assembly_—Manual_Protocol**

**1. Project Summary**

| **Item** | **Detail** |
| --- | --- |
| **Constructs** | 11 plasmids: pBEE_TrcT7_DasherGFP through pBEE_Trcp10_DasherGFP |
| **Promoters** | PRO321 (Trc-T7), PRO330–PRO339 (Trcp1–Trcp10) |
| **Strategy** | 2-fragment Gibson Assembly from single template DNA3776 |
| **Backbone PCR** | ~5,152 bp (EV_F_shared + EV_R_PROxxx) |
| **Insert PCR** | ~816 bp (insF_PROxxx + ins_R_shared) |
| **Gibson Overlap** | 34 bp complete promoter sequence |
| **Template** | DNA3776 (pBEE01_DasherGFP_1.0, 5,934 bp circular) |
| **Total Primers** | 24 (same as automated version) |
| **Total PCR Reactions** | 22 (11 backbone + 11 insert) |
| **Total Gibson Reactions** | 11 |

**2. Materials Required**

**2.1 Reagents**

| **Reagent** | **Storage** | **Amount Needed** |
| --- | --- | --- |
| Phanta Max Master Mix (Vazyme Biotech Co., Ltd) | −20°C | ~600 μL |
| Gibson Assembly 2× Master Mix, ClonExpress Ultra One Step Cloning Kit (Vazyme Biotech Co., Ltd) | −20°C | ~60 μL |
| DpnI restriction enzyme (New England Biolabs) | −20°C | ~22 μL |
| DH5α chemically competent *E. coli* | −80°C | 11 × 50 μL aliquots |
| SOC outgrowth medium | RT | ~6 mL |
| Nuclease-free water | RT | ~1 mL |
| LB agar plates + 50 μg/mL kanamycin | 4°C | 11 plates |

**2.2 Consumables**

| **Item** | **Quantity** |
| --- | --- |
| 0.2 mL PCR strip tubes or 96-well PCR plate | 22 wells |
| 1.5 mL microcentrifuge tubes | ~25 |
| Pipette tips: 10 μL, 200 μL, 1000 μL | Standard |
| PCR purification kit (magnetic beads method, Vazyme Biotech Co., Ltd) | 22 reactions |
| 1% agarose gel + TAE buffer | 1 gel |
| Ice bucket | 1 |

**2.3 Equipment**

| **Equipment** | **Notes** |
| --- | --- |
| Single-channel pipettes: P2, P10, P20, P200 |  |
| Multi-channel pipette (optional): P10, P200 | For parallel dispensing |
| Thermal cycler | Standard 96-well block |
| 42°C water bath or heat block | For heat-shock transformation |
| 37°C shaking incubator (225 rpm) | For SOC recovery |
| NanoDrop | For DNA quantification |
| Gel electrophoresis system | For PCR product QC |
| Microcentrifuge | For spin-down |

**3. Primer Reference Table**

See supplementary table 2 for full sequences.

**3.1 Shared Primers (used in all 11 constructs)**

| **Primer** | **Type** | **Length** |
| --- | --- | --- |
| **EV_F_shared** | Backbone F | 18 bp |
| **ins_R_shared** | Insert R | 50 bp |

**3.2 Construct-Specific Primers**

| **Construct** | **insF Primer (56 bp)** | **EV_R Primer (52 bp)** |
| --- | --- | --- |
| pBEE_TrcT7 (PRO321) | insF_PRO321_TrcT7 | EV_R_PRO321 |
| pBEE_Trcp1 (PRO330) | insF_PRO330_Trcp1 | EV_R_PRO330 |
| pBEE_Trcp2 (PRO331) | insF_PRO331_Trcp2 | EV_R_PRO331 |
| pBEE_Trcp3 (PRO332) | insF_PRO332_Trcp3 | EV_R_PRO332 |
| pBEE_Trcp4 (PRO333) | insF_PRO333_Trcp4 | EV_R_PRO333 |
| pBEE_Trcp5 (PRO334) | insF_PRO334_Trcp5 | EV_R_PRO334 |
| pBEE_Trcp6 (PRO335) | insF_PRO335_Trcp6 | EV_R_PRO335 |
| pBEE_Trcp7 (PRO336) | insF_PRO336_Trcp7 | EV_R_PRO336 |
| pBEE_Trcp8 (PRO337) | insF_PRO337_Trcp8 | EV_R_PRO337 |
| pBEE_Trcp9 (PRO338) | insF_PRO338_Trcp9 | EV_R_PRO338 |
| pBEE_Trcp10 (PRO339) | insF_PRO339_Trcp10 | EV_R_PRO339 |

**4. Step-by-Step Manual Protocol**

**Step 1: Primer Reconstitution & Dilution**

**Time**: ~30 min

1. Briefly centrifuge all 24 lyophilized primer tubes (12000g, 10 s).
2. Add nuclease-free water to each tube to prepare **100 μM stock**
3. Vortex 5 s, pulse-spin.
4. For each primer: transfer 10 μL stock → 90 μL water = **10 μM working solution**.
5. Label clearly and store working solutions on ice.

💡 **Tip**: Prepare primer aliquots in labeled PCR strip tubes for easy bench access.

**Step 2: PCR Setup (22 reactions)**

**Time**: ~45 min setup + ~2 h cycling

**2.1 Reaction System (50 μL per reaction)**

| **Component** | **Volume per Rxn** | **Notes** |
| --- | --- | --- |
| High-fidelity PCR 2× Master Mix | **25 μL** | Thaw on ice, vortex briefly |
| Forward primer (10 μM) | **2.5 μL** | See table below for which primer |
| Reverse primer (10 μM) | **2.5 μL** | See table below for which primer |
| Template DNA3776 (1 ng/μL) | **1 μL** | Same for all 22 reactions |
| Nuclease-free water | **19 μL** |  |
| **Total** | **50 μL** |  |

**2.2 Pipetting Order (per reaction tube)**

- **Water first**: Pipette 19 μL nuclease-free water into each tube/well.
- **Master Mix**: Add 25 μL 2X Phanta Max Master Mix (Vazyme Biotech Co., Ltd) (use P100, pipette slowly to avoid bubbles).
- **Forward primer**: Add 2.5 μL (use P2 or P10, touch wall to dispense).
- **Reverse primer**: Add 2.5 μL.
- **Template**: Add 1 μL DNA3776 (change tip between reactions to avoid cross-contamination).
- **Mix**: Pipette up/down 5× with P100 set to 25 μL, or flick tube gently.
- **Spin down**: Brief centrifuge pulse.

**2.3 Reaction Assignment Table**

| **Tube #** | **PCR Type** | **Forward Primer** | **Reverse Primer** | **Template** | **Expected Product** |
| --- | --- | --- | --- | --- | --- |
| BB-1 | Backbone | EV_F_shared | EV_R_PRO321 | DNA3776 | ~5,152 bp |
| BB-2 | Backbone | EV_F_shared | EV_R_PRO330 | DNA3776 | ~5,152 bp |
| BB-3 | Backbone | EV_F_shared | EV_R_PRO331 | DNA3776 | ~5,152 bp |
| BB-4 | Backbone | EV_F_shared | EV_R_PRO332 | DNA3776 | ~5,152 bp |
| BB-5 | Backbone | EV_F_shared | EV_R_PRO333 | DNA3776 | ~5,152 bp |
| BB-6 | Backbone | EV_F_shared | EV_R_PRO334 | DNA3776 | ~5,152 bp |
| BB-7 | Backbone | EV_F_shared | EV_R_PRO335 | DNA3776 | ~5,152 bp |
| BB-8 | Backbone | EV_F_shared | EV_R_PRO336 | DNA3776 | ~5,152 bp |
| BB-9 | Backbone | EV_F_shared | EV_R_PRO337 | DNA3776 | ~5,152 bp |
| BB-10 | Backbone | EV_F_shared | EV_R_PRO338 | DNA3776 | ~5,152 bp |
| BB-11 | Backbone | EV_F_shared | EV_R_PRO339 | DNA3776 | ~5,152 bp |
| Ins-1 | Insert | insF_PRO321_TrcT7 | ins_R_shared | DNA3776 | ~816 bp |
| Ins-2 | Insert | insF_PRO330_Trcp1 | ins_R_shared | DNA3776 | ~816 bp |
| Ins-3 | Insert | insF_PRO331_Trcp2 | ins_R_shared | DNA3776 | ~816 bp |
| Ins-4 | Insert | insF_PRO332_Trcp3 | ins_R_shared | DNA3776 | ~816 bp |
| Ins-5 | Insert | insF_PRO333_Trcp4 | ins_R_shared | DNA3776 | ~816 bp |
| Ins-6 | Insert | insF_PRO334_Trcp5 | ins_R_shared | DNA3776 | ~816 bp |
| Ins-7 | Insert | insF_PRO335_Trcp6 | ins_R_shared | DNA3776 | ~816 bp |
| Ins-8 | Insert | insF_PRO336_Trcp7 | ins_R_shared | DNA3776 | ~816 bp |
| Ins-9 | Insert | insF_PRO337_Trcp8 | ins_R_shared | DNA3776 | ~816 bp |
| Ins-10 | Insert | insF_PRO338_Trcp9 | ins_R_shared | DNA3776 | ~816 bp |
| Ins-11 | Insert | insF_PRO339_Trcp10 | ins_R_shared | DNA3776 | ~816 bp |

**2.4 Thermal Cycling Program**

| **Step** | **Temperature** | **Time** | **Cycles** |
| --- | --- | --- | --- |
| Initial denaturation | 98°C | 30 s | 1 |
| Denaturation | 98°C | 10 s | 30× |
| Annealing | 65°C | 20 s | 30× |
| Extension | 72°C | 2.5 min | 30× |
| Final extension | 72°C | 5 min | 1 |
| Hold | 4°C | ∞ | — |

**Step 3: Gel Electrophoresis QC**

**Time**: ~45 min

- Load 5 μL of each PCR reaction + 1 μL 6× loading dye onto 1% agarose gel.
- Run at 120 V for 25–30 min.
- **Expected bands**:
- Backbone (BB-1 to BB-11): **~5.1 kb** single band
- Insert (Ins-1 to Ins-11): **~800 bp** single band
- Photograph gel for records.

**Step 4: PCR Product Purification + DpnI Digestion**

**Time**: ~1.5 h

- Add **2 μL DpnI** (40 U) directly to each 50 μL PCR reaction.
- Incubate at **37°C for 1 h** to digest PCR template.
- Purify all 22 reactions using magnetic beads PCR cleanup kit following manufacturer’s protocol (Vazyme Biotech Co., Ltd).
- Elute in **50 μL** nuclease-free water.
- Measure concentration on NanoDrop.

**Step 5: Gibson Assembly (11 reactions)**

**Time**: ~15 min setup + 15 min incubation

**5.1 Reaction System (10 μL per reaction)**

| **Component** | **Volume** | **Notes** |
| --- | --- | --- |
| Purified Backbone PCR product | **2.5 μL** | BB-1 through BB-11 |
| Purified Insert PCR product | **2.5 μL** | Ins-1 through Ins-11 (matching promoter) |
| Gibson Assembly 2× Master Mix | **5 μL** | Keep on ice until use |
| **Total** | **10 μL** |  |

**5.2 Assembly Setup Table**

| **Gibson #** | **Backbone Source** | **Insert Source** | **Target Plasmid** |
| --- | --- | --- | --- |
| G-1 | BB-1 | Ins-1 | pBEE_TrcT7_DasherGFP |
| G-2 | BB-2 | Ins-2 | pBEE_Trcp1_DasherGFP |
| G-3 | BB-3 | Ins-3 | pBEE_Trcp2_DasherGFP |
| G-4 | BB-4 | Ins-4 | pBEE_Trcp3_DasherGFP |
| G-5 | BB-5 | Ins-5 | pBEE_Trcp4_DasherGFP |
| G-6 | BB-6 | Ins-6 | pBEE_Trcp5_DasherGFP |
| G-7 | BB-7 | Ins-7 | pBEE_Trcp6_DasherGFP |
| G-8 | BB-8 | Ins-8 | pBEE_Trcp7_DasherGFP |
| G-9 | BB-9 | Ins-9 | pBEE_Trcp8_DasherGFP |
| G-10 | BB-10 | Ins-10 | pBEE_Trcp9_DasherGFP |
| G-11 | BB-11 | Ins-11 | pBEE_Trcp10_DasherGFP |

**5.3 Pipetting Instructions**

- Label 11 PCR tubes (G-1 through G-11).
- Add **2.5 μL Backbone** PCR product (matching construct).
- Add **2.5 μL Insert** PCR product (matching construct). **Change tips between each!**
- Add **5 μL Gibson Assembly 2× Master Mix** (keep on ice; work quickly).
- Pipette up/down 5× gently to mix (P10, set to 8 μL).
- Spin down briefly.
- Incubate at **50°C for 15 min** in thermal cycler (lid 55°C) or water bath.
- Transfer immediately to **ice**.

💡 **Key Difference vs. Automated**: Manual pipetting of 2 μL requires extra care with P2/P10 pipette. Touch-dispense on the tube wall for accuracy.

**Step 6: E. coli Transformation (11 reactions)**

**Time**: ~2 h

- Thaw 11 × 50 μL aliquots of DH5α competent cells **on ice** (10 min).
- Add **2 μL** Gibson reaction product (G-1 through G-11) to each tube.
- **Flick tube gently 4–5 times** to mix. **Do NOT pipette up/down** — this damages cells.
- Incubate on **ice for 30 min**.
- Heat-shock at **42°C for exactly 30 seconds** (water bath preferred for uniformity).
- Return to **ice for 2 min**.
- Add **950 μL room-temperature SOC medium**.
- Incubate at **37°C, 225 rpm for 1 hour**.
- Plate **100 μL** on pre-warmed LB + kanamycin plates.
- Incubate plates inverted at **37°C overnight (14–16 h)**.
